# Control Limited Perceptual Decision Making

**DOI:** 10.1101/2022.06.24.497481

**Authors:** Juan R. Castiñeiras, Alfonso Renart

## Abstract

Periods of disengagement are generally observed during perceptual decision-making tasks, but a normative understanding of engagement is lacking. Here, we develop a theory that frames disengagement as a problem in cognitive control. Good performance through task engagement requires control, but control is costly, and this establishes a performance-control tradeoff. We derive decision policies that optimize this tradeoff as a function of the capacity of an agent for cognitive control. When their control ability is sufficiently low, agents lapse. For intermediate control limitations, a new decision-making regime appears where agents don’t lapse, but their behavior is nevertheless shaped by control. We identify hidden signatures of control-limited behavior at the level of accuracy, reaction time and decision confidence which are often observed experimentally, but had not been normatively explained. Our findings provide a path to the study of normative decision strategies in real biological agents.

---

Making the right decision often requires waiting to gather more evidence before committing to a course of action. This tradeoff between decision speed and accuracy is the essential computational problem in perceptual decision-making. Decades of experimental and theoretical work have shown that perceptual decisions are accurately described by the framework of bounded accumulation of evidence^1–5^. Many variants of this general framework have been developed and are used due to their ability to accurately describe choice, reaction time (RT) and decision confidence in psychophysical experiments^1,5–12^. In addition, these models are attractive due to their normative grounding: they describe not only how agents decide, but how they ought to decide in order to satisfy reasonable decision goals^1,13–15^. Contemporary normative theories view perceptual decision-making as a planning problem^16^ in which agents attempt to optimize payoffs in real time when the relevant state of the environment is uncertain and has to be inferred using statistical inference^17–20^.

A consistent observation in studies of decision-making is that the subject’s behavior is not always geared towards success in the task. In psychophysical experiments, this is reflected by the presence of errors far away from the subject’s psychophysical threshold^4,21–24^, which determine the asymptote of the psychometric function. These type of errors, which are referred to as lapses, are considered as a sign of disengagement. The notion of disengagement, however, is not completely clear. Whereas disengaged subjects tend to be construed as unresponsive or inattentive, strictly speaking lapses only signal behaviors which may well be proactive, but whose purpose is not aligned with the demands of the current task. Examples include exploration^23,25^, impulsivity^26,27^ or anticipation^28^, behaviors characteristic of repetitive sequential tasks, where actions are determined by events in the recent past^29–31^, or even task-engaged behaviors for a previously learned but no-longer active task^32^. In order to highlight this fact, we will refer to these as task-inappropriate behaviors.

Given that task-inappropriate behaviors compromise task-performance by definition, an analysis of their consequences is clearly relevant for normative theories of decision-making, which seek performance-optimizing policies. However, task-engagement is generally considered dichotomous, and existing normative theories apply only to engaged trials. Here, we overcome these limitations by framing the management of task-inappropriate behaviors as a problem of cognitive control. In our framework, a perceptual decision-making task implicitly defines a behavioral policy which is optimal from the point of view of task performance, but which is costly from the point of view of cognitive control. Indeed, efficiently trading off speed and accuracy requires both suppressing actions until enough information has been obtained^26,28,33^ and also attention^34–36^, both of which are triggers of cognitive control^27,37–40^. Task-inappropriate policies, on the other hand, are costly in terms of performance, but we reason that they are observed because they represent existing strategies available to the agent which are cheap in terms of control. Thus, we propose that a normative description of decision-making behavior should describe a tradeoff between maximization of task performance and minimization of the amount of cognitive control needed to overcome task-inappropriate behaviors.

Here, we develop a mathematical framework that describes the optimal tradeoff between control and performance costs in perceptual decision-making, as a function of the capacity for cognitive control of an agent. Fully engaged, or stimulus-independent task-inappropriate policies are limiting cases in this tradeoff, but for a range of intermediate levels of cognitive control, novel decision regimes appear which display signatures of accuracy, RT and decision confidence observed but previously unaccounted-for within a normative framework.

## Optimal perceptual decision-making

We first introduce optimal perceptual decision-making as it’s typically studied, in the absence of control limitations. We model the structure of a typical decision-making experiment in the laboratory, with one binary decision per trial (comprising choice, RT and decision confidence), difficulty varying randomly across trials, a time penalty for errors, and an inter-trial interval (Fig. 1a). To formalize the optimal strategy in this task, the sensory stimulus is described by the analog value of a scalar sensory feature (constant within a trial). The stimulus is not directly observed, but the agent has access to noisy ‘observations’, in which the stimulus is corrupted by temporally uncorrelated zero-mean noisy samples. In general, the stimulus value is drawn from a symmetric distribution centered around zero on each trial, and the task of the agent is to decide whether the stimulus belongs to the positive or negative category. This requires inferring the sign of the latent stimulus value on each trial based on the stochastic observations, and reporting it though a categorical binary choice^18^ (Fig. 1b top). The absolute magnitude of the latent stimulus defines the strength of evidence and thus the trial’s difficulty. The agent begins the trial undecided and with the correct prior over the value of the latent state, and uses the observations to sequentially update the posterior probability over the stimulus (Fig. 1b middle). Since the task only requires a report on the sign of the latent state, all future consequences depend only on the agent’s running belief that the latent state is positive, which we denote by *g*(*t*) and which we will simply refer to as ‘belief’. This recursive inferential process defines a stochastic trajectory on *g*(*t*) in each trial (Fig. 1b, bottom).

**Fig 1.**
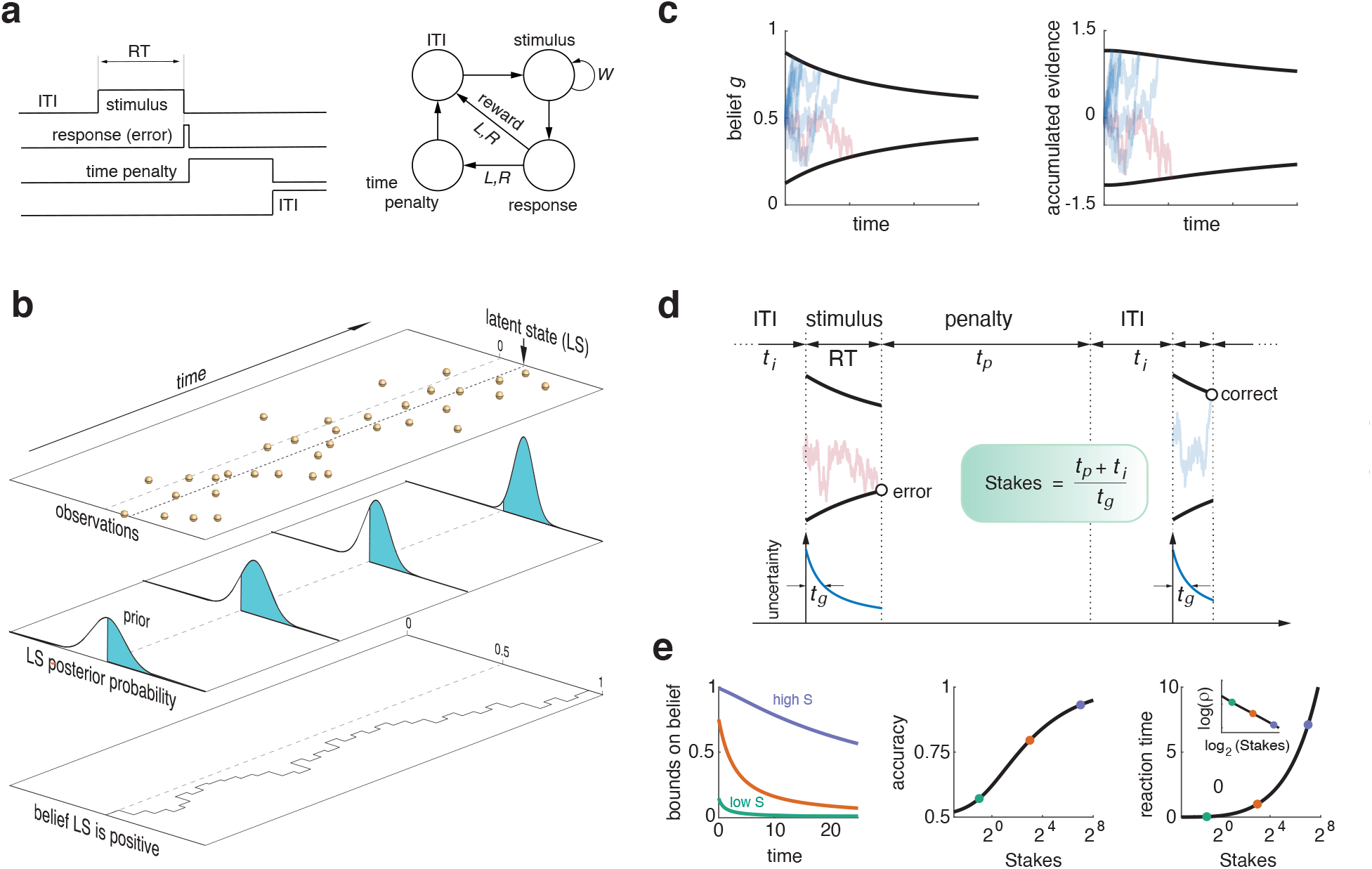
Optimal perceptual decision-making without control limitations. **(a)** Trial-structure of the tasks we study. Left. Time sequence of events during an example (error) trial. Right. State space with possible transitions in the task. *W* stands for wait. **(b)** Schematic description of the inferential process in one trial of a sequential sampling problem. The agent experiences noisy observations whose mean (black dotted line) defines the analog, latent, value of the stimulus (top). The task of the agent is to decide if the latent state is positive or negative. To do this, it first updates its prior belief about the value of the latent state based on the incoming information, into a posterior distribution. As more information is sampled throughout the trial, the mean of the posterior approaches the true mean and the posterior uncertainty decreases (middle). Outcomes depend only on the belief that the latent state is positive (the area under the posterior on the positive side; bottom), which fluctuates stochastically driven by the noisy observations. Observations come at discrete time-steps in this example for illustration purposes, but time is continuous in our model. **(c)** Optimal decision bounds on belief (Left) and on accumulated evidence (Right) from a set of trials. Note the one-to-one map between belief and accumulated evidence at every moment. **(d)** Definition of the task Stakes. The panel shows a series of trials during a behavioral session. During the stimulus, the uncertainty of the agent about the latent state decreases. The timescale of inference is the time it takes for the agent’s uncertainty to decrease by a factor of two (Supplementary Information). The task Stakes is the sum of the penalty time and inter-trial interval in units of the timescale of inference. **(e)** Left. Decision bounds on belief for three values of the task Stakes. Middle, Accuracy averaged across all difficulties (equal to the decision confidence^41^) as a function of the task Stakes. Right. Same for RT. Inset. Reward rate decreases monotonically with the Stakes (inset).

Because trials are typically short, but there are many of them in one session, we assume that the goal of the agent is to maximize the ‘reward rate’ across the whole session (see Supplementary Information for details on how to use this performance measure). This allows the description of agents sensitive to the long-term consequences of their actions without the need to invoke temporal discounting^42,43^. Reward rate (which we denote by the symbol *ρ*) is the standard performance measure in foraging theory^44^ but has also been used in perceptual decision-making^1,18,45^.

Using the theory of Dynamic Programming^46^, previous studies showed that the reward rate maximizing policy is for the agent to commit to either choice when its instantaneous belief *g*(*t*) reaches a certain time-dependent decaying bound^17,18^ (Fig. 1c, left; Supplementary Fig. 1). Because the stochastic sensory evidence is temporally uncorrelated, there is a one-to-one monotonic map between belief and accumulated evidence, which depends on elapsed time during the trial^18^ (Supplementary Information). Thus, the time-dependent decision bound on belief is equivalent to a different time-dependent bound on accumulated evidence (Fig. 1c right), implying that the optimal policy has a neural implementation in terms of a drift-diffusion model^17,18,47^, as long as the temporal evolution of the decision bounds can be specified accurately.

How is the speed-accuracy tradeoff (SAT) resolved by the optimal agent? Interestingly, it can be shown (Supplementary Information) that a single effective parameter spans the universe of optimal decision-making policies and, in particular, arbitrates the SAT (Supplementary Fig. 2). This parameter, which we have termed the task Stakes, is equal to the sum of the penalty time and inter-trial interval, measured in units of the timescale of inference (Fig. 1d). As the agent samples the stimulus, its initial uncertainty about the value of the stimulus decreases. We designate the timescale of inference *t*_*g*_ as the time it takes for the initial stimulus uncertainty to become one half of its initial value (Supplementary Information). Intuitively, the Stakes measures whether trials are frequent or infrequent, relative to the approximate time it takes to discriminate the stimulus. If trials are very frequent in relative terms, then it does not pay off for the subject to invest time in performing well in the task, since another opportunity for reward with at least 0.5 probability will be presented immediately. In this case we say the Stakes for the agent are low, and the optimal policy entails a bound on belief close to 0.5 which is associated to uncertain decisions regardless of stimulus difficulty. On the other extreme, if trials are very rare in relative terms, then the agent ought to invest time to try to obtain reward in these rare opportunities, i.e., the Stakes for the agent are high. In this case, the bound on belief is high and trials tend to involve high-confidence decisions (Fig. 1e, left). As expected, when the Stakes are raised, both accuracy and RT increase (Fig. 1e middle, right), implying that the Stakes control the resolution of the SAT by the optimal agent. As the Stakes grow, RT increases faster than accuracy, and the effective time between trials also increases, leading to a net reduction in reward rate (Fig. 1e right, inset). Experimental manipulations of the ‘opportunity cost of time’ during perceptual decision-making induce similar relationships between reward rate and the SAT^48^. We typically vary the time penalty *t*_*p*_ to set the task Stakes. This leads to the intuitively reasonable scenario where, as the time penalty increases, the overall reward rate of the agent goes down.

## The performance-control tradeoff and optimal control-limited policies

We developed a normative strategy to describe the effect of task-inappropriate behaviors in decisionmaking by framing it as a problem in cognitive control. To do this, we assume that the agent possesses a default policy *P*_def_(*a*) (Fig. 2a). The possible actions *a* during the stimulus are *L* and *R* (which imply commitment to either a Left or a Right response – we define the *R* response to be correct when the stimulus is positive) and *W* (Wait), which implies postponing commitment to a later time. Because the default policy describes a task-inappropriate behavior, the agent under this policy makes stimulus-independent binary choices which look random from the point of view of the sensory discrimination task (i.e., lapses). This assumed randomness is purely phenomenological, and simply summarizes all task-independent factors which can shape the agent’s responding. We usually assume unbiased *L* and *R* response probabilities (Fig 2a left), but default responses can be biased (Fig 2a right) and can depend on events in the previous trial, which we use to model sequential dependencies. Importantly, the default policy also interferes with the task-adaptive timing of action by specifying a default rate of responding, which we term *λ* (i.e., under this policy, the mean RT = *λ*^−1^). Thus, *λ* determines the agent’s default responsiveness (Fig. 2a left) and biases the resolution of the SAT away from the optimum defined by the task Stakes (Fig. 1d-e). When *λ* is large, the agent is impulsive and thus biased towards speed. When *λ* is small, the agent is unresponsive and thus biased against speed, or towards accuracy (simply because it accumulates evidence for a longer time). Although more complex default policies can be considered, these capture the essential consequences of task-inappropriate behaviors from the point of view of the decision-making task, while remaining mathematically tractable.

**Fig. 2.**
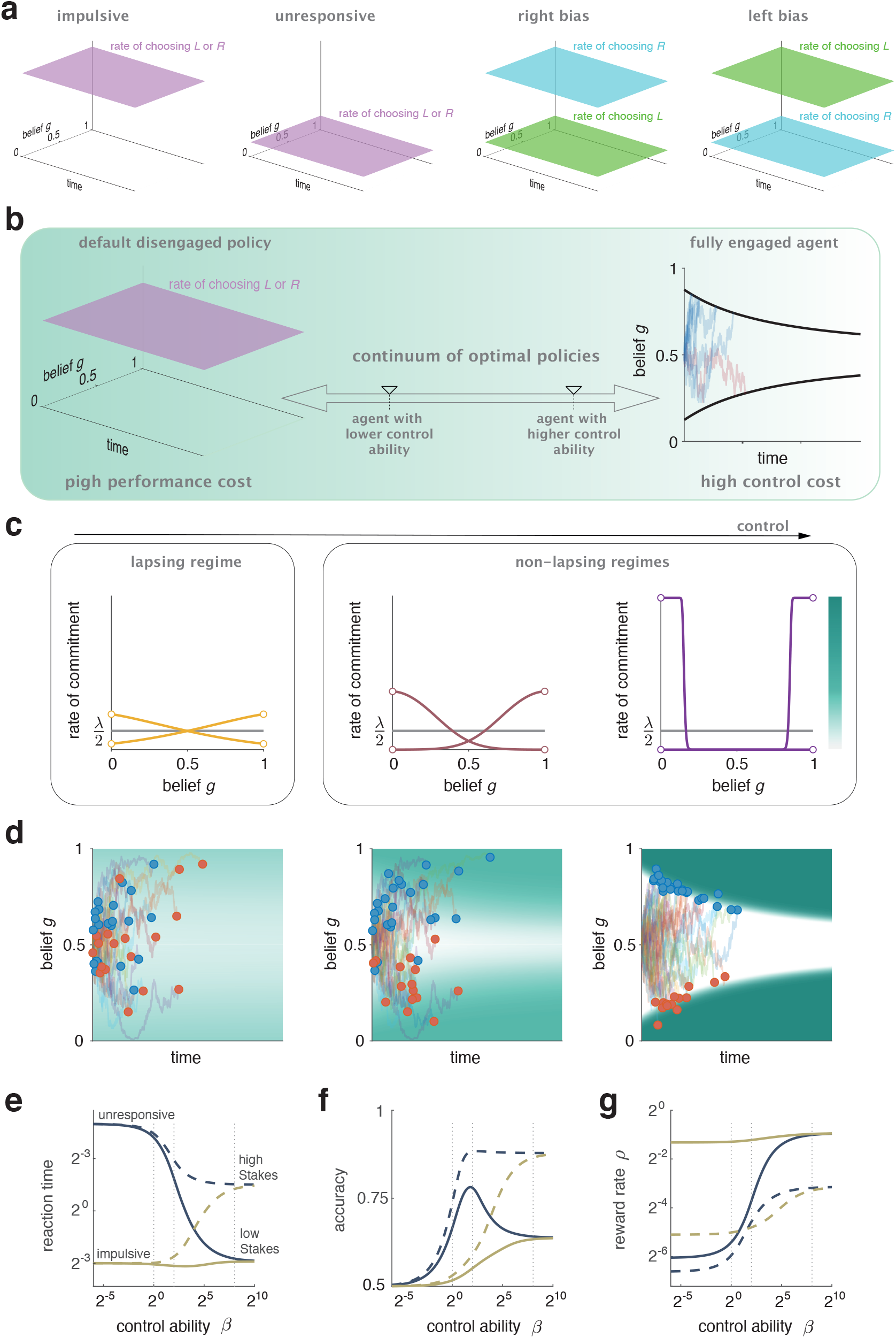
Modeling control-limited decision-making. **(a)** Default policies we consider in our study, represented as instantaneous probabilities per unit time of commitment. Default policies are stimulus-independent, i.e, they don’t depend on the agent’s belief *g*. The left two panels correspond to random *L*/*R* choices with different degrees of impulsivity. The right two panels describe *R* and *L* biases. **(b)** Schematic description of the tradeoff between performance and control. **(c)** Optimal control-limited policies at a given time during stimulus presentation, for agents capable of increasing control (left to right). The horizontal gray line in all panels is the default rate of commitment. **(d)** Belief trajectories for multiple trials for the three agents displayed in (c). In each trial, the agent’s belief evolves stochastically driven by the evidence, and commitment occurs according to the optimal policy. The instantaneous rate of commitment in each case is color-coded in the plot background, according to the colorbar besides panel (c), right. **(e)** RT (averaged across difficulties) as a function of the control ability of the agent *β*, for two different levels of responsiveness (color) and two different penalty times *t*_*p*_ (implying different task Stakes; solid versus dashed lines). The three control regimes in (c,d) are shown as vertical dashed lines **(f**,**g)** Same for Accuracy and for the reward rate *ρ*. Parameter values for results in all figures are displayed in Supplementary Table 1.

The default, or automatic, status that we give to the task-inappropriate policy formally means that we assume there are no costs in terms of cognitive control incurred by the agent when using it. However, because default choices are stimulus-independent, there is a large performance cost associated to the default behavior. On the other hand, the optimal policy described in the last section has the least possible performance costs, but requires mobilizing cognitive resources, which are costly^49–52^. This situation defines a tradeoff between performance and control (Fig. 2b). To formalize this tradeoff, one can enlarge the total cost minimized by the agent so that it includes both performance and control terms. Namely, we seek an optimal policy *P*_opt_(*a*|*s*) that minimizes a total cost

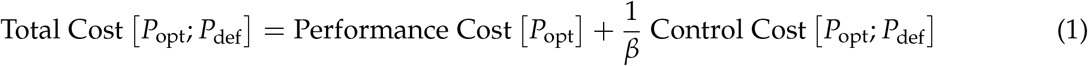

The optimal policy does depend on the ‘environmental states’ *s* which, as we described in the last section, are the belief *g* and the elapsed time *t* since the beginning of the trial (Supplementary Information). For the control cost, we use the Kullback-Leibler (KL) control framework^53–55^, in which the cost of control is equal to the KL divergence *KL [P*_opt_|*P*_def_] between the optimal and default policies – a quantitative measure of the dissimilarity between two probability distributions^56^ (Supplementary Information). The performance cost is the negative expected future reward of the agent, i.e., the standard objective function in a sequential decision problem. Although we formulate the problem here in terms of costs, a formulation in terms of ‘rewards’ (negative costs) is totally equivalent (Supplementary Information).

The control aspect of the problem has thus two elements. First, the term *KL [P*_opt_|*P*_def_] measures the amount of conflict that the task induces for the agent, in line with frameworks where conflict drives the use of cognitive control^57^. If the default response tendencies are aligned with the current task demands, it will be close to zero, whereas it will be large if, in a given state, the task requires a different action from the one the agent would choose by default. Second, the relevance of this kind of conflict in shaping the behavior of the agent is given by the positive constant *β*^−1^, which determines the relative importance of performance and control costs. When *β*^−1^ is large, deviating from the default policy is very costly, so we use this constant to quantify the control limitations of the agent at a given moment.

To obtain optimal decision policies for control-limited agents, we used techniques from optimal control theory to minimize Eq. 1 with respect to *P*_opt_(*a*|*s*). As before, we actually minimize the ‘total cost rate’ across a behavioral session (Supplementary Information; for parsimony, we continue to refer to this as maximizing reward rate despite the fact that it includes the cost of control). The optimal policies take the form of instantaneous probabilities of commitment per unit time to the *L* or *R* responses as a function of belief and time, and can thus be construed as a stochastic ‘race’ model^1^ in which the agent’s decision is to the response that commits first. Let’s consider the probability per unit time of commitment to, say, the *R* response, at a given moment *t* within the stimulus presentation. As a function of belief, this probability will be constant and equal to *λ*/2 for agents incapable of exerting any control (*β* = 0), whereas for agents without control limitations (*β*^−1^ = 0) it will be zero for beliefs *g* < *B*_*u*_(*t*) and infinite for beliefs *g* ≥ *B*_*u*_(*t*), where *B*_*u*_(*t*) is the value of the upper optimal decision bound at time *t* (Fig. 1 c). For intermediate values of *β*, control-limited agents display sigmoidal commitment rates which smoothly interpolate between these two extreme situations (Fig. 2c).

Examination of these policies permits the definition of three different regimes. For sufficiently low control, neither the probabilities of *R* or *L* commitment are zero even under complete certainty about the nature of the sensory stimulus (i.e., for *g* = 0, 1; Fig. 2c left). In this regime, the psychometric function of the agent will display lapses, because the lower probability (incorrect) response can still be chosen first, even under absolute certainty about the stimulus category. This reflects the fact that, for this agent, task-relevant information has a very limited control over behavior. For sufficiently high control, the optimal policies look like step functions which resemble hard decision bounds. In this regime the *L* and *R* commitment policies are never active simultaneously (Fig. 2c right) and the default policy has effectively no influence over behavior during the task. For intermediate levels of control, a third regime exists. In this regime, the probability of an incorrect response when the agent is certain is zero, so the agent does not lapse. But when the agent is uncertain, the *L* and *R* commitment policies are both active and compete (Fig. 2c, middle). This represents a novel decision-making regime because while, as we show below, accuracy, RT and decision confidence are shaped by the imperfect suppression of task-independent random choices, these effects are ‘hidden’, and cannot be identified from the presence of decision lapses.

The probabilistic nature of the control-limited policies can be characterized as involving ‘smooth’ decision bounds, which permit commitment over broad ranges of belief at any given time (Fig. 2d). To understand the consequences of limited control for decision-making behavior, we first looked at the aggregate (across difficulties) accuracy and RT of the agent. For increasing control, RT (Fig. 2e) and accuracy (Fig. 2f) transition from being fully determined by the default policy to being fully determined by the task Stakes. The reward rate always increases monotonically with the control ability of the agent (Fig. 2g), establishing a relationship between motivation and control, which can be potentially exploited by the agent. Examining the behavior of the control-limited agents as a function of the Stakes (Supplementary Fig. 3) reveals more clearly that they are poorly adaptable, and only perform well when the task demands are aligned with their default response tendencies.

## Decision lapses in control-limited agents

As we have just noted, if control limitations are sufficiently strong, the psychometric function of the optimal agent will display lapses (Fig. 3a). From the point of view of psychophysics, the relevance of lapses is that they obscure the true psychophysical performance of the subject. Because perceptual decision-making experiments are used to probe the limits of sensory discrimination accuracy, developing methods to recover true sensory limitations in the presence of non-sensory behavioral processes has been a critical problem in the history of psychophysics. For instance, Signal Detection Theory arose in the 1950s from the realization that false alarms are not guesses^58^. The standard approach to recover a ‘clean’ estimate of stimulus discriminability in the presence of lapses is simply to scale vertically the psychometric function until its asymptotes reach one and zero^21^. This is appropriate when engagement is viewed as a dichotomous all-or-none process, because the resulting ‘standard’ correction is equivalent to inferring, and then removing, the fraction of trials where choice is assumed to be completely random – which is determined by the asymptote of the psychometric function (Fig. 3b). In contrast, when lapses result from control limitations, which are not dichotomous, the proper approach to recover the true sensory limitations of the agent is to examine the psychometric function at *β*^−1^ = 0, i.e., in the absence of any limitations in control.

**Fig. 3.**
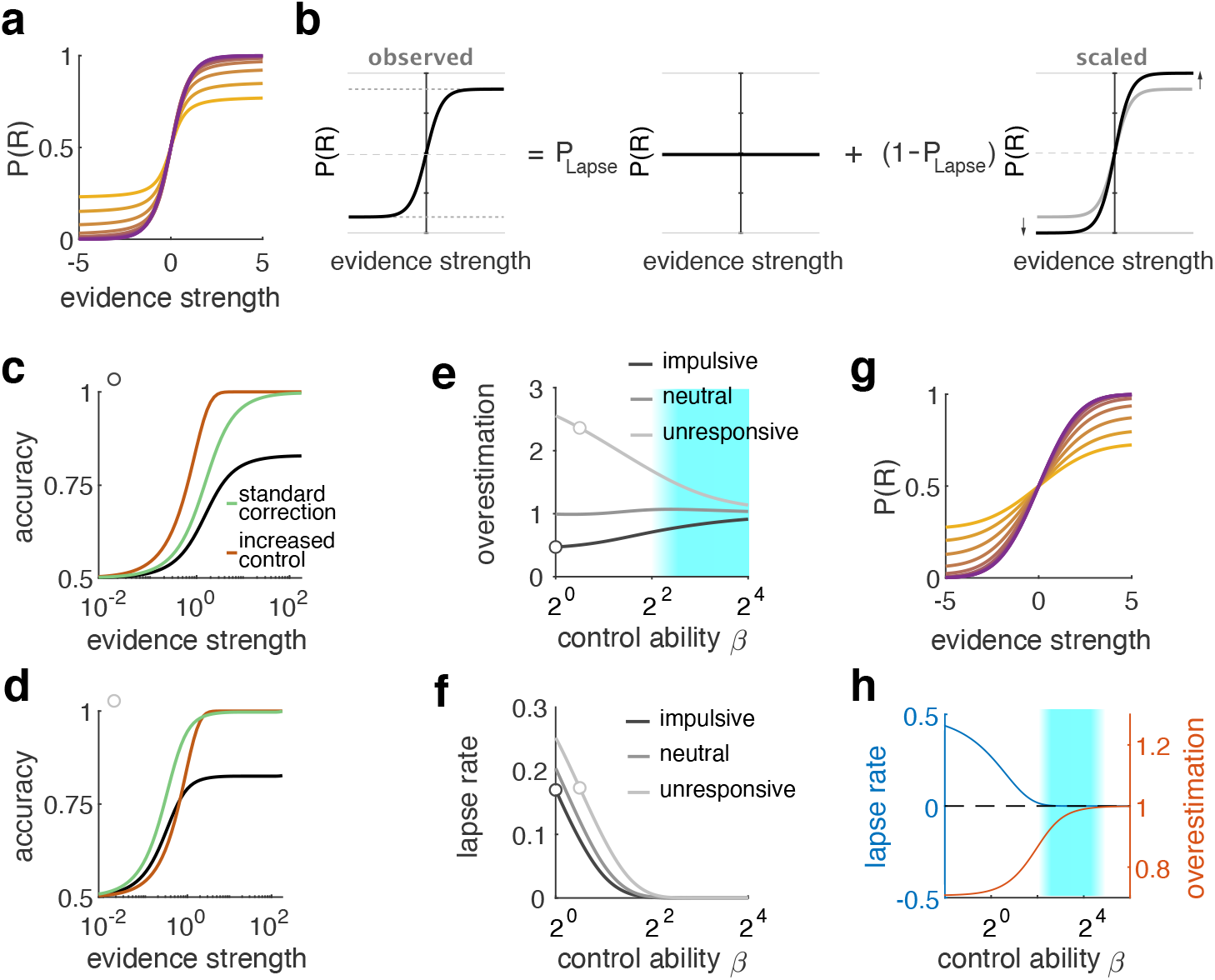
Decision lapses produced by control limitations. **(a)** Psychometric functions display increasing lapse rates as the control-limitations of the agent grow (orange to blue). **(b)** Standard strategy for recovering the true discrimination ability of the agent in the presence of lapses by scaling the observed psychometric function. **(c)** Psychometric function for an impulsive (speed-biased) control-limited agent (black) in the lapsing regime. Green is the standard correction (equal to the black curve scaled) Red is the correction obtained by setting *β*^−1^ = 0 (the psychometric function of the agent without control limitations for the same value of the Stakes). **(d)** Same as (d) but for an accuracy-biased agent. **(e)** Ratio between the slope of the psychometric function corrected with the standard method and the psychometric function of the agent without control limitations, as a function of the control ability *β*, for three levels of responsiveness. The two examples in (c,d) are signaled with a circle. When the ratio is larger than 1, the standard correction overestimates the discrimination accuracy of the subject. The cyan background, here and in (h) is the novel regime in Fig. 2c, middle. **(f)** Lapse rate as a function of *β* for the same three levels of responsiveness as in (e). **(g)** Psychometric functions in a ‘single sensory sample’ model^23^ (equivalent to a SDT setting) as a function of the control ability of the agent. **(h)** Lapse rate (blue) and degree of overestimation by the standard correction (same as (e)) for this model.

To compare these two approaches, we considered the result of applying both types of corrections to the psychometric function of a control-limited agent. Interestingly, the two approaches don’t, in general, coincide (Fig. 3c,d). In fact, the ‘cleaned’ psychometric slope obtained using the standard correction can either over- or under-estimate the slope of the psychometric function of the optimal agent without control limitations, depending on the default response rate *λ* (Fig. 3e). When the lapsing agent is impulsive, control-limitations are detrimental for performance, and the standard correction underestimates the discrimination ability of the agent (Fig. 3c). Conversely, when the lapsing agent is too slow, larger reward rates can be achieved by emphasizing speed at the expense of accuracy, in which case the standard correction overestimates discriminability (Fig. 3d).

When we examine how lapse rates depend on the control ability *β* (Fig. 3f), we see that the discrepancy in the slope of the corrected psychometric functions (Fig. 3e) persists after the agent is no longer lapsing. Thus, in this regime (corresponding to the intermediate control situation in Fig. 2c middle, and signaled by the cyan background in Fig. 3e), control limitations shape discrimination accuracy but are hidden (because lapses are not observed), concealing the true psychophysical ability of the agent. This phenomenon is still present when one considers a simpler decision problem where an agent only receives one sample of sensory evidence (equivalent to a SDT setting). Here, one can also define optimal control-limited policies which will display lapses if the control ability of the subject is sufficiently low^23^ (Fig 3g). In this setting, the psychometric function does not depend on speed-accuracy considerations, and the standard correction factor always underestimates the true psychometric slope (Fig. 3h). These results suggest that the absence of decision lapses is not an accurate reporter of fully engaged behavior.

## Sequential dependencies and decision biases

In laboratory settings, where many decisions are performed during an experiment, it is often observed that behavior in one trial can be partly explained by events taking place in previous trials^30,59–62^. Sequential dependencies might interact with control in different ways depending on whether they are part of the default policy (as would be the case, for instance, if the agent made a decision in the current trial based on the outcome of their previous choice regardless of the sensory stimulus) or not. In this last case, they could reflect a fully adaptive task strategy (for instance if the environment itself contains sequential correlations^30^), or a partially adaptive strategy in the face of processing limitations (as when agents sequentially update the decision boundary in the task^61,62^). Often, many of these sources of sequential dependence operate simultaneously^30^.

We consider the interaction between control and three classes of sequential dependencies, grouped according to the quantity that is updated from one trial to the next. One class corresponds to updating the predisposition of choosing an action before the stimulus is observed, which can be modelled using biased default response policies (Fig. 4a-c, Fig. 2a right). Another class corresponds to updating the value of the different actions, depending on previous events (Fig. 4d-f). A third class corresponds to updating the prior belief of the agent about the upcoming stimulus category (Fig. 4g-i). We devised a procedure for deriving optimal policies that incorporate each of these three forms of trial-to-trial updating (Supplementary Information). Importantly, each class can be used to describe a number of qualitatively different sources of sequential dependence. For instance, updates in the default probability of choosing an action can depend on the previous choice, or on an interaction between the previous choice and outcome. Our grouping into classes thus reflects the target of the cross-trial updating, not the events that cause the update. To reveal the effects of these different forms of sequential dependence, we plot the psychometric functions of the agent conditioned on the relevant event in the previous trial. We use as a baseline an agent whose control limitations lead to a substantial lapse rate, because in this regime the effect of different forms of sequential dependence are more clearly distinguishable. Then we show how these effects are modified as the agent becomes fully control-capable.

**Fig. 4.**
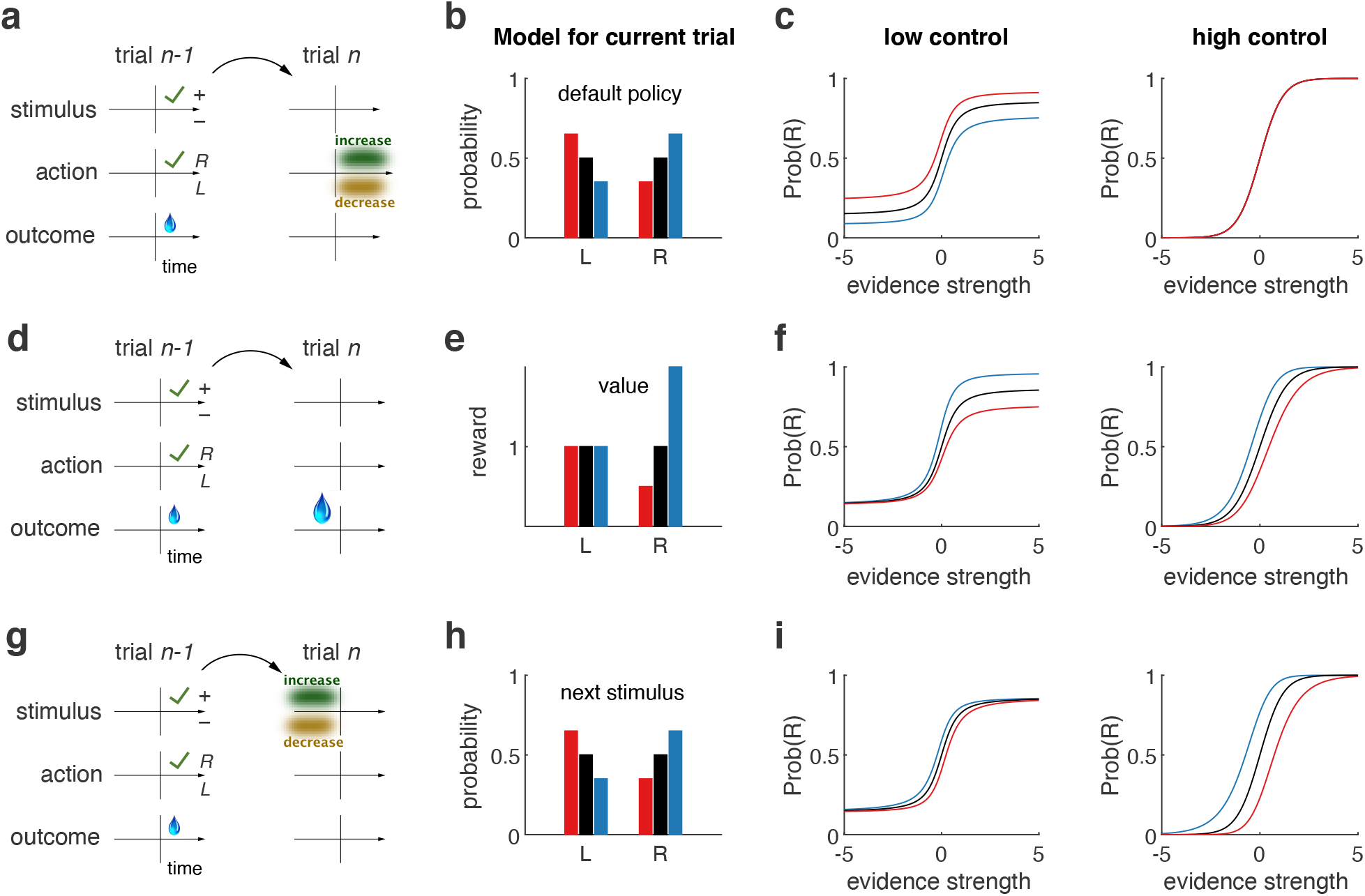
Interaction between sequential dependencies and control. **(a)** Schematic description of sequential dependencies. Left and right columns show stimulus sign, action and outcome for the previous and current trial respectively. The vertical line in each plot marks the stimulus onset. Here, events in trial *n* − 1 increase (decrease) the default probability of responding *R* (*L*) in trial *n*. **(b)** Three scenarios where the default policy is unbiased (black) or biased towards action *R* (blue, as in (a)) or *L* (red; Methods). **(c)** Psychometric functions for each of the three cases in (b) for control-limited (left) and control-capable (right) agents. **(d)** Similar to (a), but where trial history shapes the current value of the action that the agent just performed. **(e)** Three scenarios where the agent choose *R* in the previous trial and the history of choice-outcomes experienced leads to the reward magnitude in the current trial being modelled as higher (blue, as in (d)), lower (red) or average (black). **(f)** Same as (c) for value biases. **(g)** Similar to (a) but where trial history shapes the agent’s prior beliefs about the upcoming stimulus, depicted as a shading over the stimulus categories before the vertical bar marking stimulus onset. **(h)** Three scenarios where the agent believes the latent variable in the current trial is more likely to be positive (blue, as in (g)), negative (red) or is unbiased (black). **(i)** Same as (c) but for biases in stimulus probability.

The signature of sequentially updating the bias of the default policy, is a symmetric vertical displacement in the psychometric function, as observed, for instance, by Scott et al.^63^ (Fig. 4c, left). This is because increasing the probability of one action automatically implies lowering the probability of the other. These vertical shifts are unoccluded when the psychometric function displays significant lapsing. Because sequential biases are only present in the default policy, they reflect a control-limitation and disappear for control-capable agents (Fig. 4c, right).

Agents might, on the other hand, use their history of successes and failures to sequentially update the value of each action (Fig. 4d), instead of the default probability of choosing it. In experiments where rewards and penalties are fixed, this would be a form of suboptimality, but it might be expected if subjects have the wrong model for the task, and incorrectly attribute fluctuations in average value across trials (due to variable proportions of incorrect choices) to fluctuations in the single-trial value of an action^64^. Because only the value of the action that was just produced is updated (Fig. 4e), this type of sequential dependencies lead to asymmetric modulations of the psychometric function, in which the amount of bias is proportional to the likelihood of repeating the action (Fig. 4f, left), as was recently observed^23^. In this case, although the sequential bias is still there for control capable agents, the marked asymmetry is almost completely eliminated (Fig. 4e, right), because the probability of repeating the action is already saturated at its maximum value of 1 when lapses disappear.

A final scenario we consider is an update in the prior belief of the agent about the stimulus (Fig. 4g-h) which might arise, for instance, if there are across-trial correlations in the value of the latent variable which the agent is learning^30^. Updating of stimulus priors is expected to lead to horizontal displacements in the psychometric function. In a sequential sampling framework there is an approximately horizontal displacement (Fig. 4i) but, since in this case the sequential bias is fully adaptive and the default policy is unbiased, the horizontal shifts in the psychometric function are more salient in the control-capable agent.

These results show that the shape of the modifications in the psychometric function due to trial history, together with their dependency on the agent’s control ability, can be used to infer which aspect of the decision-making process is being updated across trials.

## Control limitations shape decision confidence and reaction time

The smoothing of the decision bounds caused by the control-limitations of the agent (Fig. 2d) strongly shapes the transformation from sensory evidence into categorical choice, leading to characteristic patterns of RT and decision confidence (DC). DC in a categorical choice measures the decision-maker’s belief in her choice being correct^65–69^. Explicit judgements (or implicit measures^70^) of DC depend on various properties of the decision problem, such as discrimination difficulty, trial outcome or time pressure^65,71–73^. We compared patterns of DC in control-capable (regime in 2c right) and control-limited (regime in 2c middle) agents during performance of easy and hard discriminations. In both discriminations, the latent variable is positive (i.e., *R* is the correct choice), but we varied the strength of the evidence. We recall that *g*(*t*) describes the agent’s belief that the latent variable is positive. Thus, DC is given by the value of *g*(*t*) at the moment of commitment for rightward decisions, and by 1 − *g*(*t*) for leftward ones. When the strength of evidence is weak, individual trials are compatible with beliefs spanning both choice options depending on the stochastic evidence (Fig. 5a left), whereas when the strength of evidence is large, the agent’s beliefs quickly converge on rightward preferences (Fig. 5a right).

**Fig. 5.**
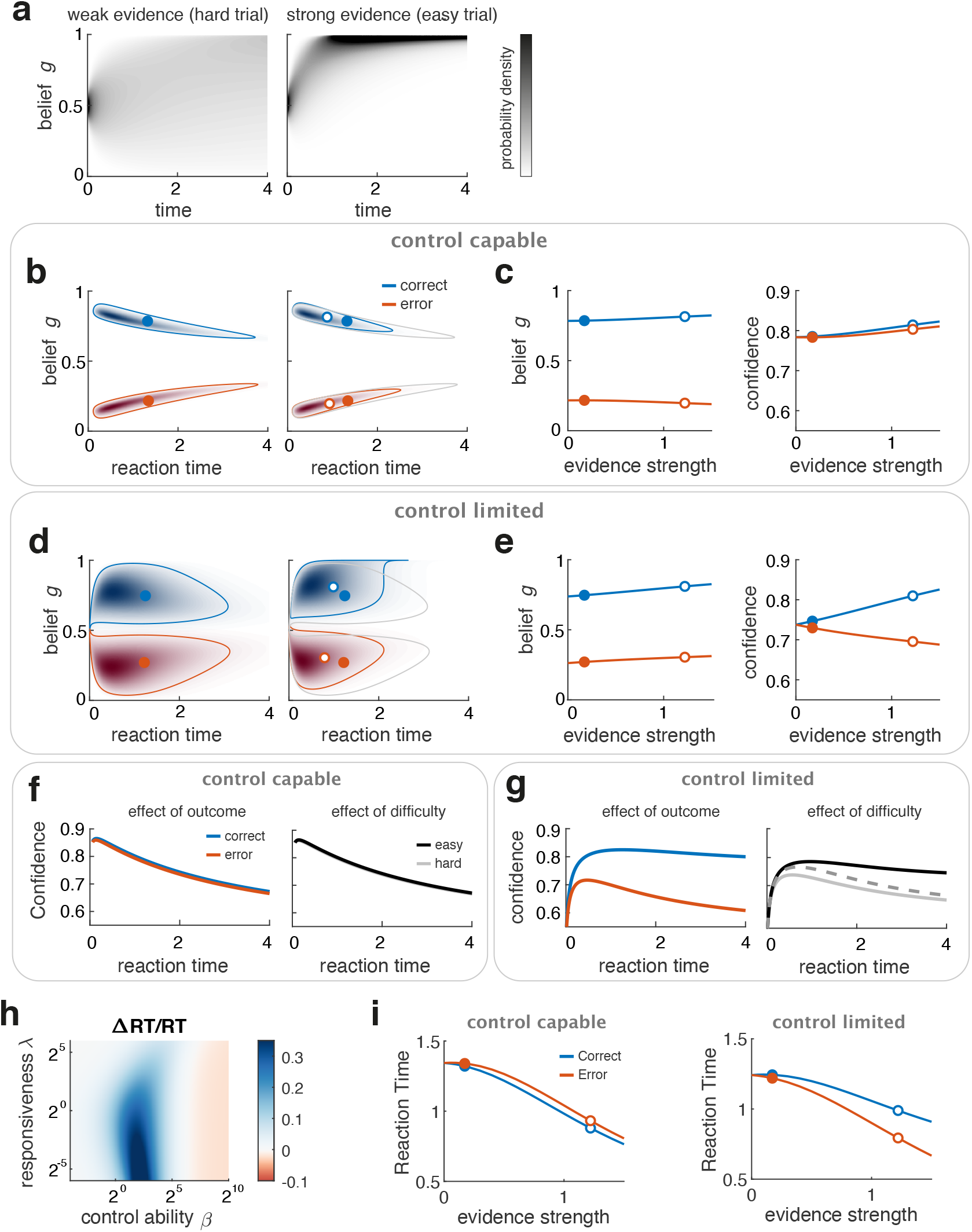
Control limitations shape decision confidence and reaction time. **(a)** Temporal evolution of belief for a difficult (left) and easy (right) stimulus condition. The probability distribution of beliefs (Methods) is normalized at each time. **(b)** Left. Joint distribution of belief (at decision time) and RT for correct and error trials for the difficult condition (a, left) for an agent without control limitations (as in Fig. 2c, right). Circles represent the mean. Contour demarcates the region enclosing 90% of the probability mass. Right. Same but for the easy condition in (a, right). Mean is now shown as a white circle. The means and regions of high probability from the hard condition are also shown for comparison. **(c)** Left. Mean belief at decision time as a function of the strength of evidence for correct and error trials. Circles show the values used in (a,b). Right. Decision confidence (DC, equal to 1 − *g* if *g* < 0.5) as a function of the strength of evidence. **(d**,**e)** Same as (b,c) but for an agent in the moderate control regime (Fig. 2c middle). **(f)** DC conditioned on RT as a function of outcome (left) and difficulty (right) for the control-capable agent in (b,c). **(g)** Same as (f) for the moderately control-limited agent in (d,e). In the right panel, we also show the tachometric function (accuracy as a function of RT; dashed). **(h)** Relative difference in RT as a function of outcome as a function of the control ability *β* and the default rate of responding *λ*. **(i)** Chronometric curves for correct and error trials for control capable (left) and moderately control limited (right) agents.

In order to more accurately quantify the behavior of our agents, we developed methods to compute semi-analytically the joint distribution of DC and RT corresponding to any of the optimal policies (Supplementary Information). If control limitations are negligible, this joint distribution is tightly focused (Fig. 5b, see also Fig. 2d right) around the regions where commitment abruptly transitions to a non-zero value (Fig. 2c right), approximating a temporally decaying hard bound^18^ (Fig. 1c left). When the strength of evidence grows, the distribution still tracks the symmetric bounds but is shifted towards earlier RTs, and thus more extreme beliefs (Fig. 5b, right; Fig. 5c, left). Because the bounds are necessarily outcome-symmetric, DC is almost outcome-independent and grows with the strength of evidence (Fig. 5c, right). The outcome-dependence of DC is referred to as confidence resolution (CR)^65,71,72^. Interestingly, the optimal decision policy without control limitations, which is always superior in terms of reward rate, has poor CR^20^. A tendency for decision confidence to increase with the strength of evidence regardless of outcome is sometimes observed^67,74,75^, specially in experiments requiring simultaneous reports of decision confidence and choice^67,74^.

For the control-limited agent, the joint distribution of DC and RT is much less concentrated (Fig. 5d; see also Fig. 2c middle). When the strength of evidence is almost zero, the distributions for correct and error trials are still approximately symmetric (Fig. 5d, left). But for easy conditions both distributions shift up towards rightward beliefs (i.e., towards the evidence), making them asymmetric with respect of outcome (Fig. 5d, right). Intuitively, this occurs because the control-limited policy, despite being outcome-symmetric, is less restrictive in terms of the values of belief and RT where commitment is possible, so when beliefs are strongly biased by the evidence, DC ends up also reflecting this bias (Fig. 5d, right, Fig. 5e, left). Since errors occur when the agent believes *L* is the correct choice (*g* < 0.5), the biasing of these beliefs by the evidence (towards *R*, i.e., towards *g* = 1) implies a more undecided state (*g* closer to 0.5), i.e., lower DC. This process results in opposing trends for DC as a function of the strength of evidence for correct and error trials (Fig 5e, right), demonstrating that optimal control-limited policies possess good CR. CR is generally observed in psychophysical experiments^12,68,73,76–79^, but had so far been unaccounted within a normative sequential sampling framework. Control-capable agents have poor CR because, when choices are triggered by hard bounds on belief, the only quantity that can shape DC is RT (through the time-dependence of the decision bounds). In fact, if the bounds were constant, as in the SPRT, DC would be identical for all choices, as noted early on^80^. Thus, for agents using these kinds of policies, DC conditional on RT is independent of any other aspect of the problem, such as trial outcome or strength of evidence (Fig. 5f). In contrast, DC in control-limited agents is shaped by all factors that affect the beliefs of the agent before commitment, and is therefore larger for correct than error trials, and for easier compared to than harder conditions (Fig. 5g).

Figs. 5f,g reveal that the time-course of accuracy and DC vs RT depends on control (note that, averaged across stimulus difficulty, accuracy and DC are equal^41^). The curve displaying accuracy conditional on RT is called the tachometric function. The optimal control-capable policy has a monotonically decreasing tachometric function^67^ (Fig. 5f). In contrast, tachometric functions for control-limited policies display an initial gradual rise (Fig. 5g). The initial rise takes place because very short RTs are necessarily associated to uncertain beliefs.

The outcome-dependence of RT is also affected by limitations in control. Experimentally, the outcome-dependence of RT has been shown to depend on various factors such as stimulus difficulty or the speed-accuracy regime^8,12,80–83^. Efforts to match the observed patterns of outcome-dependence has lead to developments in decision-making theories^6,8,84,85^, but so far these have largely relied on mechanistic considerations. The difference between mean RT for correct and error trials accurately signals the control-regime of the agent (Fig. 5h). For agents whose behavior is not limited by control limitations, errors are slightly slower than correct trials (Fig. 5i, left). This occurs because the decision bounds decrease with time (Supplementary Fig. 4 describes the relationship between the outcome-dependence of RT and the slope of the decision bounds). Conversely, in the novel control-limited regime, errors are faster than correct trials (compare Fig. 5h and Fig. 3e,h). Errors tend to occur earlier because, on the one hand, the stochastic control-limited policy allows commitment with uncertain beliefs and, on the other hand, these beliefs are more likely earlier on, since the belief of the agent aligns with the evidence as the trial progresses (Fig. 5a,d; see also Supplementary Fig. 5 for an explanation using a minimal version of the stochastic policies).

While the results in Fig. 5 show that a single underlying factor (variations in the amount of control exercised by the agent) can span the different regimes of outcome-dependence in DC and RT observed experimentally, we have not yet discussed the factors that might shape the control ability of an agent. We turn to this question next.

## Control allocation and task regimes

So far, we have assumed that the control ability is a stable property of an agent. While there are stable differences in cognitive control across different species^86^ (in decision-making tasks, for instance, human subjects generally tend to lapse less than rodents^4^), the ability to exercise control is both a dynamic and a limited resource, specially in human subjects^52,87–89^. Furthermore, this limited resource can be allocated strategically, taking into account the gains that might be experienced from different allocation policies^90,91^. Although a theory of optimal trial-by-trial control allocation is beyond the scope of the current study, we sought to understand what gains can be expected from the allocation of control under different task regimes.

We focus on speed-accuracy and task-difficulty manipulations, which are commonly used. Task difficulty is reduced by lowering the timescale of inference parameter *t*_*g*_ (Fig. 1d), whereas speed emphasis is achieved by lowering the task Stakes by reducing the penalty time *t*_*p*_ and inter-trial-interval *t*_*i*_ (Fig. 1d-e), while keeping *t*_*g*_ fixed. Although changing the timescale of inference *t*_*g*_ also modifies the Stakes (Fig. 1d), the two types of manipulations are not equivalent, because changing *t*_*g*_ also modifies the units of time of the problem, which directly affects the reward rate (see Supplementary Information). Figs. 6a,b show how the reward rate of the agent depends on the control ability *β* and on the task Stakes, or the task difficulty, respectively. When *β* is fixed, the reward rate decreases with the task Stakes (Fig. 6a, Fig. 1e, right) and also with increasing difficulty (Fig. 6b).

**Fig. 6.**
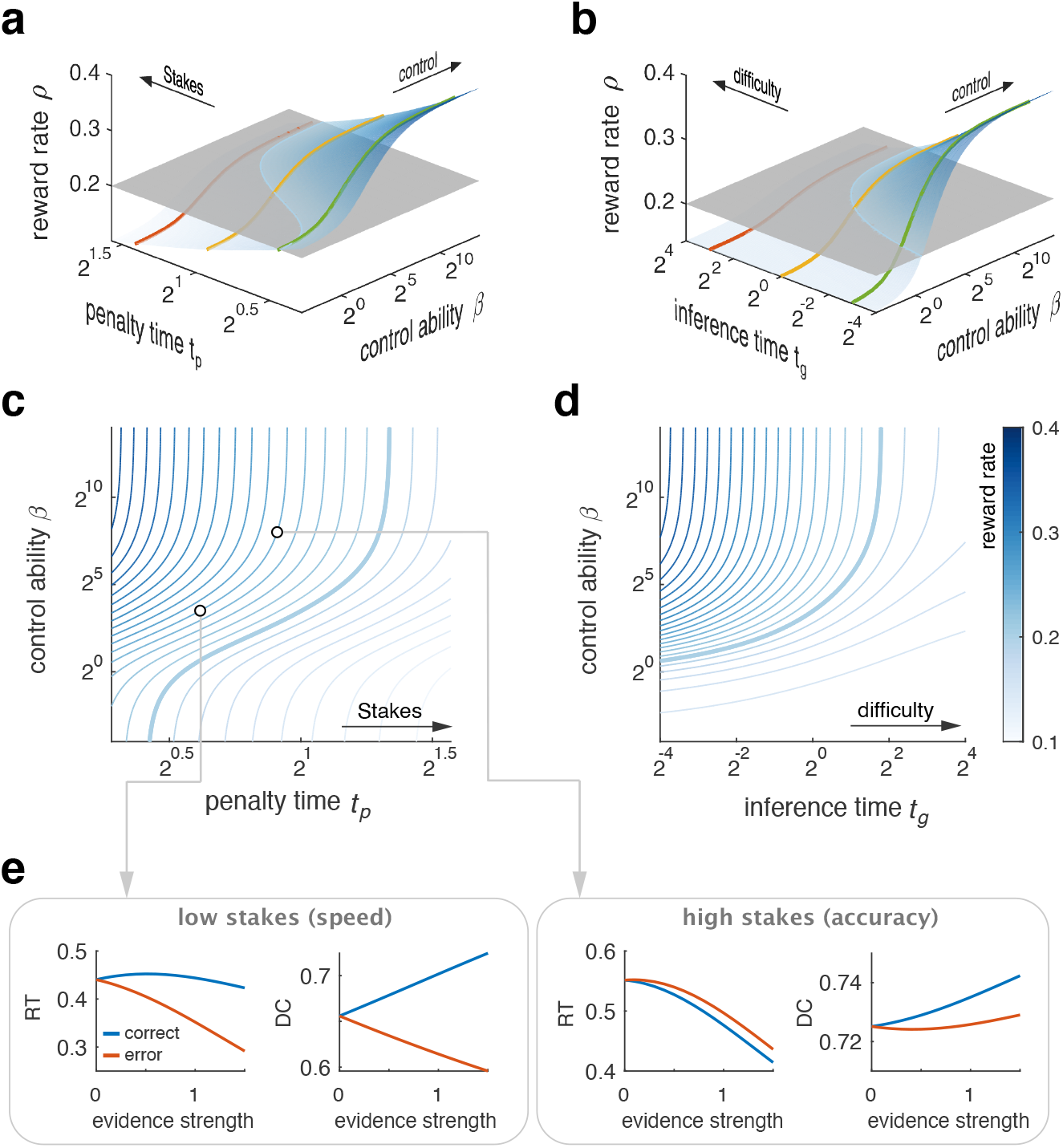
Efficient allocation of control across task regimes explains outcome dependence of RT and DC. **(a)** Reward rate as a function of control ability *β* for three values of the task Stakes (controlled through the penalty time *t*_*p*_). Curved surface spans all values of *t*_*p*_. The intersection between this surface and each constant reward-rate plane defines an implicit map between the task Stakes and the amount of control that needs to be allocated to reach that reward rate. **(b)** Same procedure to define an implicit map between control and task difficulty. **(c)** Implicit maps between control and the task Stakes for different target reward rates. Circles represent values of *β* and *t*_*p*_ in the examples in (e). Thin line represents the map shown in (a) **(d)** Same as (c) but for the map between control and difficulty. **(e)** Left. Outcome-dependence of RT and DC for a low-Stakes task – where speed is normatively required – at the level of control needed to achieve a particular reward rate. The resulting policy has fast errors and clear confidence resolution. Right. When the task Stakes are raised – which demands accuracy – more control is needed to reach the same reward rate. The corresponding policy has slow errors and low confidence resolution.

In addition, as mentioned previously, for every task setting, the reward rate of the agent always grows with the control ability *β*. Thus, if the agent’s goal was to maximize reward rate under any conceivable circumstance, higher invested control would always pay off. However, it is more plausible that agents seek to obtain higher reward rates only until the experienced reward rate is sufficient, a form of satisficing^45,92,93^. We therefore postulate that, given the limited capacity for exercising control^52,87–89^, agents will, for each task setting, use the minimal amount of control necessary to reach their reward target^94^. This mechanism induces an implicit map between task regimes and control investment, for any given reward target. The implicit map can be revealed by examining the constant reward rate surfaces as a function of the control ability *β* and parameters defining the task regime, such as *t*_*p*_ (Fig. 6a) or *t*_*g*_ (Fig. 6b). This analysis shows that, for every reward target, the implicit map induces a positive covariation between control and accuracy emphasis (Fig. 6c) or task difficulty (Fig. 6d).

These results suggest that normative agents should tend to display the signatures of moderate control policies when the task difficulty is low, or when speed is emphasized because, for any reward target, in these conditions the target can be achieved with a lower investment of control (Fig. 6e). This qualitative pattern is exactly what the psychophysical literature reveals: For easy discriminations or under time pressure, errors tend to be faster than correct trials^8,12,81,82^ and subjects display higher confidence resolution^12,78^, whereas difficult discriminations and accuracy emphasis are associated with slower errors^8,12,80,82,83^ (Fig. 6e). Also consistent with this trend, decision lapses are more frequently observed in primate psychophysical tasks during the initial training stages, where task difficulty is low^95^. Thus, adaptive patterns of control allocation provide a normative explanation for how the properties of a discrimination task shape the outcome-dependence of RT and DC and the presence of decision lapses.

## Discussion

We have described how to optimally trade-off control and performance costs in perceptual decisionmaking, and have argued that this represents an attractive normative framework to describe the problem of task engagement relative to task-inappropriate behaviors. Our work thus endorses the view that an essential aspect of behavior is the strategic arbitration of different ‘behavioral controllers’, which are adaptive in different contexts or over different timescales, or are sensitive to different environmental cues^33,96,97^. We contend that, for certain tasks such as perceptual decisions, there is a clear difference between these controllers in terms of the amount of cognitive resources that they require. In this sense, the essence of task-inappropriate behaviors as we construe them is that they require minimal cognitive control, and thus represent default modes of behavior which the agent will fall back onto despite the performance cost that they entail.

It might be argued that task-inappropriate behaviors are observed because animals have a hard-wired drive to explore, not because these behaviors are less cognitively costly. Exploration itself, however, is under the influence of control. On the one hand, this is supported by the observation that task-inappropriate behaviors are less prevalent in human subjects^4^, which have evolved more sophisticated control faculties^86^. Some studies have addressed explicitly the interplay between the exploration-exploitation tradeoff and control using *n*-armed bandit tasks^98,99^. It was initially suggested that, in fact, it is exploitation that should be considered as an automatic default, and that exploratory choices require cognitive control^98^. For instance, behavior driven by Pavlovian associations might be construed as automatic. However, other studies have found that, even in bandit tasks, cognitive load (which limits the ability to use cognitive control) makes behavior more stochastic^100,101^. A recent study directly probed the relationship between cognitive load and directed versus random exploration, and found that only directed exploration is down-regulated in high-load conditions^102^. Besides, it is critical to note that exploitative behavior in an *n*-armed bandit task is very different from what might constitute exploitative behavior in a perceptual decision-making task, which requires attention^35,36^ and withholding responses during temporal accumulation of evidence^27,39,40^.

The performance-control tradeoff also has implications for the study of motivation, as the control ability of the agent can only be defined relative to the reward available from the different actions (i.e., *β*^−1^ has units of reward – Supplementary Information). Motivation has been construed as a map that specifies the utility of outcomes^103^, which is posited to vary across different internal states. Control-limitations provide another dimension that affects the efficiency with which outcomes can mobilize behavior. The same reward will be differently effective in driving adaptive responding, even under equal internal states, depending on how such adaptive responses relate to an agent’s default policies and its capability for cognitive control.

An important novel aspect of our framework is that it does not automatically tie suboptimal engagement with the presence of decision lapses. This is because, unlike existing frameworks, we don’t model engagement as dichotomous, but rather as a graded process dependent on the allocation of control. When the control costs of staying engaged are sufficiently high, agents indeed lapse (Fig. 3a), but when control-costs are moderate, no lapses are visible in the psychometric function. Nevertheless, we’ve shown that the phenomenology of decision-making in this regime is fundamentally control-limited (Figs. 4,5). The reason moderately control-limited agents don’t lapse is that we model control demands as arising from the presence of conflict^57^. A stimulus-independent policy and the optimal decision policy conflict the most when the sensory evidence is largest. Thus, conflict is reduced most effectively by shaping these easy conditions first. In practice, the consequence of this fact is that many of the signatures of partial engagement are more subtle than the presence of lapses in the psychometric function – which have been the focus of existing theories^23,24^ – and are evident at the level of reaction time, decision confidence, the tachometric function, and the shape of the psychometric curve.

Lapses arising from control limitations are formally similar to lapses construed as a form of exploration, as recently suggested by Pisupati et al.^23^ in the context of a Signal Detection Theory framework. However, whereas we view lapses as essentially reflecting a limitation of the agent, Pisupati et al. construe them as adaptive in perceptual decision-making tasks – because the agent will perceive action-outcome as being stochastic due to errors caused by their sensory limitations. We think it’s unlikely that the existence of sensory errors *per se* will generally be modelled by subjects, including rodents, as reflecting probabilistic reward contingencies, given that rats in some difficult perceptual decision-making tasks do not lapse^5^. However, exploratory strategies would be adaptive^99^ if the agent learns an incorrect model of the task, as in this case the agent will construe the environment as probabilistic.

Optimal control-limited policies take the form of a stochastic race model. How should this stochastic commitment rules be understood? Although our model is not mechanistic, it could be approximated by a model using a conventional deterministic bound but in the presence of a compound decision variable depending on both the accumulated sensory evidence (which determines the agent’s instantaneous belief, Fig. 1c; Supplementary Information), and a stimulus-independent stochastic term, which would induce variability in the transformation from accumulated evidence (and thus belief) to action. The control ability of the agent would be reflected in the relative weight of these two inputs to the net decision variable. A related family of models with compound decision variables has recently been proposed^28,104^, which involve a race between two stimulus-dependent and -independent components, and where the stimulus-independent component depends on elapsed time. The presence of a stimulus-independent stochastic term effectively implements a smooth commitment rule and, as such, these models produce initially rising tachometric functions^28,104^, and fast errors^104^. We suggest that this phenomenology can be understood as an adaptive response to a limitation in control. Indeed, the stimulus-independent term in the rats of Hernandez et al.^28^ implements anticipation to the onset of the stimulus^5^, which leads to premature responding and is thus costly in terms of performance and reward rate. Such ‘Pavlovian’ anticipatory behaviors are often task-inappropriate^97^. An even coarser approximation to our stochastic policies would be to consider models with a deterministic decision bound within a trial, but which varies randomly across trials^8^. It terms of control, however, frameworks based on suppression of inputs (embodying task-inappropriate policies) to action selection circuits seem more parsimonious. We have analyzed in detail how control-limitations shape the phenomenology of perceptual decision-making tasks. A salient feature of control-limited agents is that they display strong confidence resolution (CR; Fig. 5d) in contrast to the control-capable policy (Fig. 5c). The standard mechanistic solution to obtain robust CR within a sequential sampling setting relies on post-decisional processing of DC^12,73,78,79,105^. Our results show that stochastic commitment is another source of CR which, additionally, has a natural normative interpretation. Furthermore, although control-limited policies and post-decisional processing both produce CR robustly, they are clearly distinct, and result in opposing trends in terms of the outcome-dependence of RT and of the tachometric function, quantities which are unmodified by post-decisional processing. We have argued that the observed reversal of the outcome-dependence of RT as a function of task difficulty or speed-accuracy regime, can be understood in terms of strategic control allocation. Mechanistically, a related, but distinct, manipulation would be to use a faster stimulus-independent component in models with a compound decision variable^104^. Whereas this seems natural in response to an increased demand for speed, it’s not clear why it would be an adaptive response to a decrease in difficulty. It will be interesting to examine how the outcome dependence of RT varies with respect to explicit control manipulations, such as the presence of concurrent tasks^35^

Our model is an extension of the KL control framework^53–55,106^. Initially developed as providing efficient solutions to optimal control problems in engineering, this framework is also finding applications in behavioral neuroscience, such as tractable solutions to the flexible-replanning problem^107^, or as an explanatory framework for capacity limitations in action-policy complexity^55,101^. From a methodological perspective, our main contribution is a flexible mathematical framework for optimal control of non-stationary point processes in uncertain environments (Supplementary Information). Here, these model the timing of discrete actions, but the framework can be naturally extended to model spiking neural activity, an approach we are currently pursuing. An important elaboration of our theory will be to go beyond describing limitations in the control of action and incorporate explicit limitations in the process of inference itself^108^. Even more critical will be to model explicitly the dynamics of control allocation. Although partial solutions to this problem have been proposed^91^, they don’t involve planning (in the sense of incorporating the long-term consequences of action) nor sensory uncertainty. We studied strategic control allocation assuming agents use satisficing strategies to obtain ‘good enough’ reward rates (Fig. 6). Interestingly, both satisficing and KL control itself can be considered ways of making resource-limited or bounded rational decisions^92^ – in the case of KL control, in the sense of finding optimal policies which are good enough in terms of complexity^109^. Extensions of our framework addressing the trial-by-trial allocation of control would provide a normative approach to study the existence of persistent engaged and task-inappropriate behaviors across sessions^24^.

## Methods

Analytical and numerical methods are described in the Supplementary Information.

## Code Availability

Custom MATLAB scripts used to implement the mathematical framework and produce the figures are available upon request.

## Acknowledgements

We thank Pietro Vertecchi, Tiago Costa, and Gautam Agarwal for discussions, and Davide Reato, Caroline Heimerl, Zach Mainen, Daniel McNamee, Joe Paton, Christian Machens, and Jaime de la Rocha for comments on the manuscript. J.C. was supported by a doctoral fellowship from the Fundação para a Ciência e a Tecnologia (FCT). AR was supported by the Champalimaud Foundation, a Marie Curie Career Integration Grant PCIG11-GA-2012-322339, the HFSP Young Investigator Award RGY0089, the EU FP7 grant ICT-2011-9-600925 (NeuroSeeker), and grants LISBOA-01-0145-FEDER-032077 and PTDC/MED-NEU/4584/2021 from the FCT.

## Author contributions

J.C and A.R. conceived the project and the theory. J.C developed the theory and conducted the analysis. A.R. wrote the manuscript with feedback from all authors.

## Supplementary Figures

**Supplementary Fig 1.**
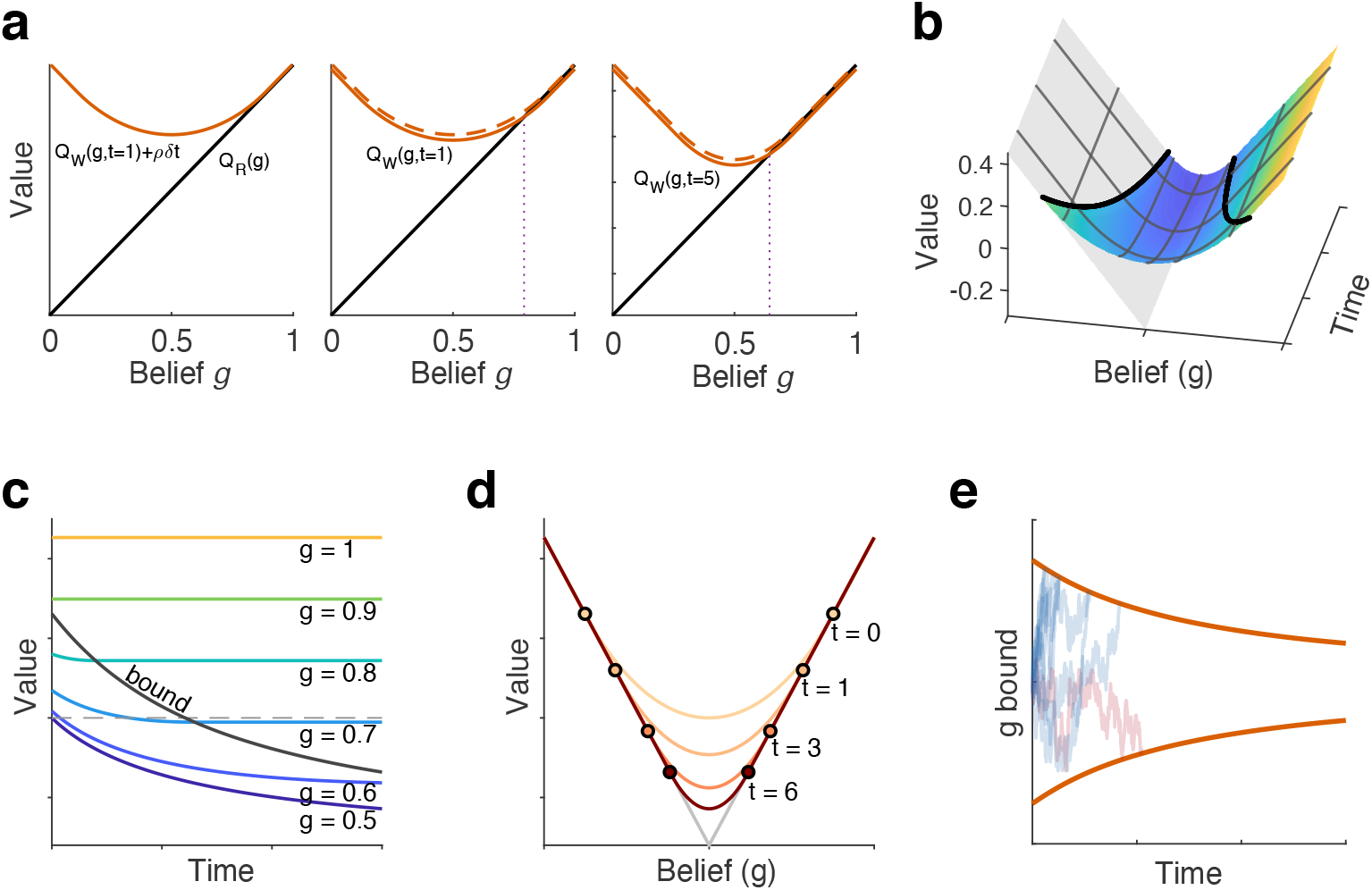
Optimal perceptual decision-making using dynamic programming. **(a)** Procedure for obtaining the optimal bounds on belief. Left. Action-value of the *R* choice (black) as a function of belief *g*, and action-value of waiting *W* without considering the opportunity cost of time *ρδ dt* (orange). The orange curve is tangent to the black one, as the information gained by waiting cannot decrease the Value. Middle. When one considers the opportunity cost of time (new solid line, previous curve is now dashed), the action-values of choosing *R* and *W* intersect, defining the optimal decision bound on belief at this time (*t* = 1). Right. Same procedure, but for a later time (*t* = 5), shows that the decision bounds decrease with time. **(b)** The full value function as a function of belief and time, with the decision bounds overlaid in black. Action value of choosing right is shown as a gray plane, to illustrate the intersection with the value function. Notice how the two functions are tangent over the decision bounds marking the intersection. Several constant-belief and constant-time curves are also shown. **(c)** The value function for fixed beliefs as a function of time. For any belief lower than the decision-bound at *t* = 0, the value of that belief initially decreases and then becomes constant (when it crosses the bound). The value of uncertain beliefs decreases initially because the increase in belief from every subsequent increment of evidence decreases with time (Supplementary Information). **(d)** Value function for different times. Circles indicate the location of the decision bound at that time. All curves are convex and symmetric with respect to *g* = 0.5. The action value of choosing right and left is also shown in gray to illustrate again that the curves are tangent above the bound. **(d)** Examples of belief trajectories up to the bound in a few trials.

**Supplementary Fig 2.**
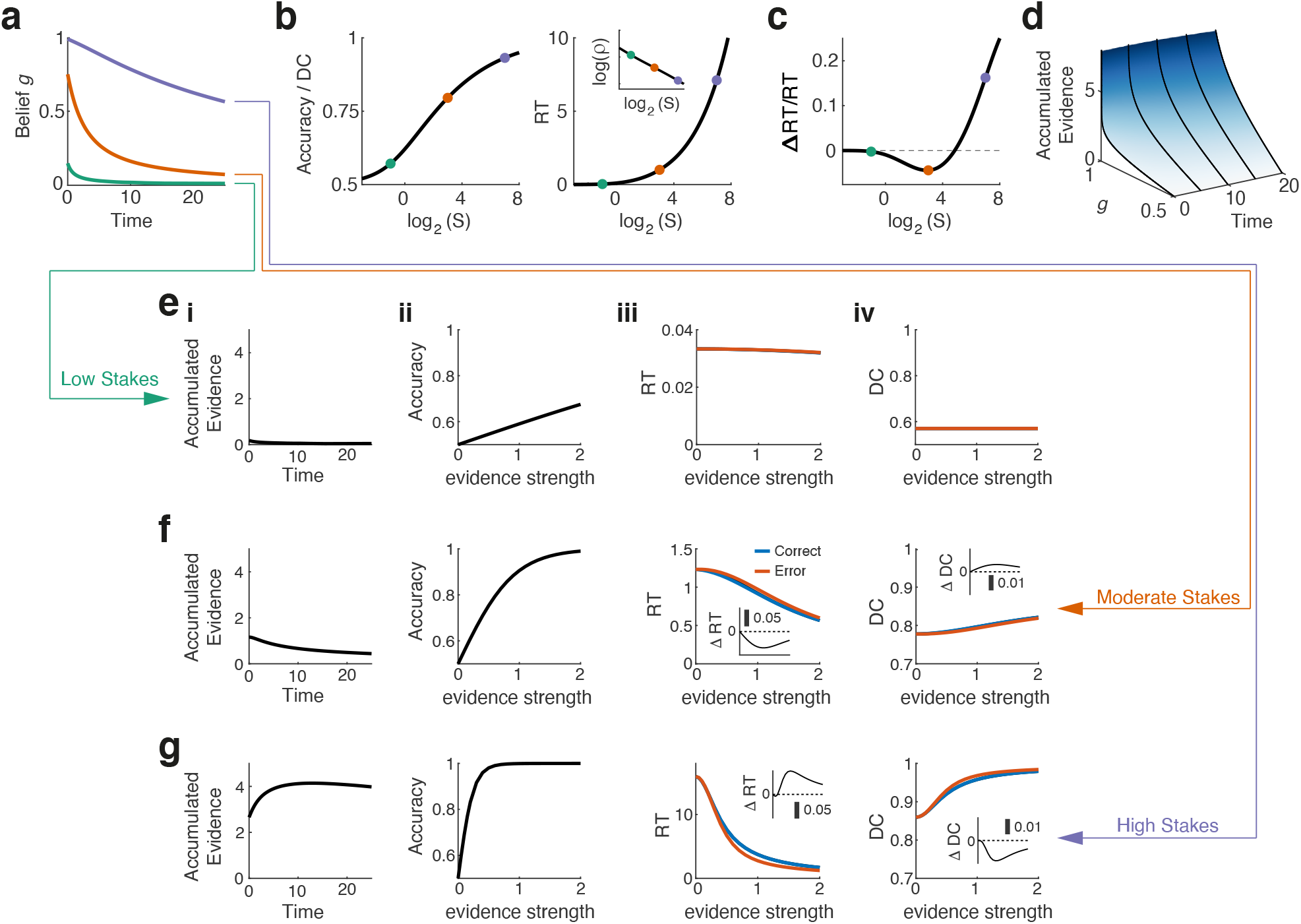
Effect of the task Stakes on optimal decision policies for agents without control limitations. **(a)** Decision bounds on belief for three values of the Stakes. **(b)** Accuracy averaged across all difficulties (equal to the decision confidence; Left) and RT (Right) as a function of the Stakes. Inset. Reward rate decreases monotonically with the Stakes. Panels (a-b) as in Fig. 2. **(c)** Relative difference in (average) RT between correct and error trials as a function of the Stakes. Notice that this quantity is non-monotonic and achieves a negative minimum for the intermediate value of the Stakes in (a). The range of Stakes for which the quantity is negative corresponds to the typical range used experimentally. **(d)** One to one map between instantaneous belief *g* and time, and accumulated evidence during the trial. Although the bounds on belief always decay with time (a), extremely large values of accumulated evidence are necessary to reach large values of *g*(*t*), specially at long times, resulting in a situation where the optimal bounds on accumulated evidence initially grow with time when the Stakes are sufficiently high (f *i*). This ultimately determines the reversal of the sign of ∆RT/RT (see also Supplementary Figure 4). **(e)** Behavior of the agent for a low-Stakes task. *i*. Decision bound on accumulated evidence. *ii*. Psychometric function. *iii*. Chronometric functions for correct (blue) and error (red) trials. *iv*. Decision confidence as a function of evidence strength for both outcomes. In panels (*ii iv*) the upper limit on the strength of evidence is twice the width of the prior. **(f**,**g)** Same as (e) but for moderate and high Stakes respectively. Insets in (*iii, iv*) show the difference in RT and Decision confidence, respectively, between correct and error trials. Notice the anti-correlation between ∆RT and ∆DC, consequence of the decaying bound in belief, and also the sign reversal between (f) and (g), consistent with (b).

**Supplementary Fig 3.**
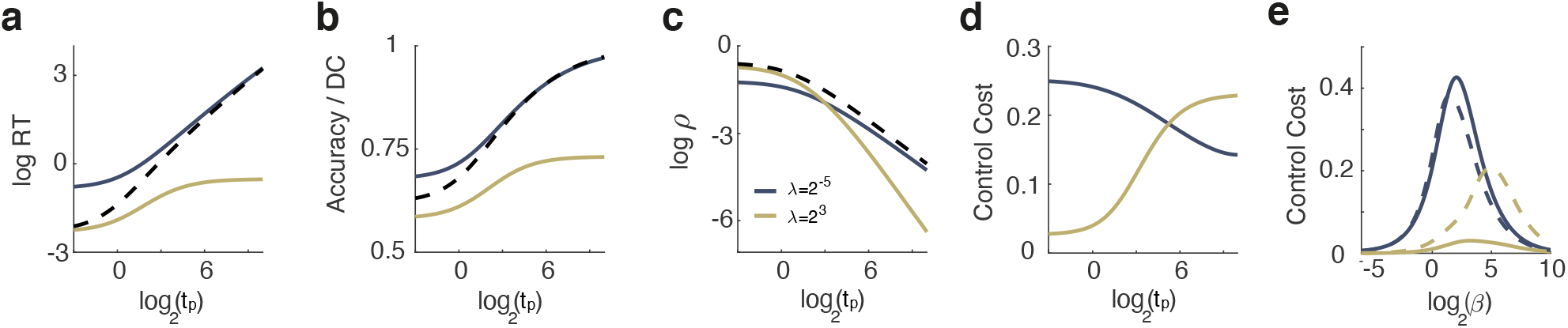
Effect of the task Stakes on the behavior of control-limited agents. **(a-d)** RT, Accuracy/DC, reward rate and control cost (*β*^−1^*KL*[*P*_opt_ | *P*_def_]) respectively, as a function of the task Stakes, for two moderately control-limited agent (as in Fig. 2c middle, in this example *β* = 2^4^) with different default policies (different colors). For comparison, in (a-d) the black dashed line shows the behavior of the agent without control limitations. Control-limited agents don’t adapt their behavior efficiently to the demands of the task (as represented by the task Stakes), so their behavior approximately matches that of the control capable agent only when the task demands are roughly aligned with their default tendencies. **(e)** Control cost (as in (d)) as a function of the control ability of the agent. The control cost is always zero for extreme values of the control ability. When *β* is close to zero, control is too costly for the agent, so the optimal strategy is to operate under the default policy, in which case the *KL* term vanishes. At the other extreme, when *β*^−1^ is near zero, the control cost is zero because there are no control limitations. Throughout this figure, we fix *t*_*g*_ = *t*_*i*_ = 1, so that the Stakes = *t*_*p*_ + 1.

**Supplementary Fig 4.**
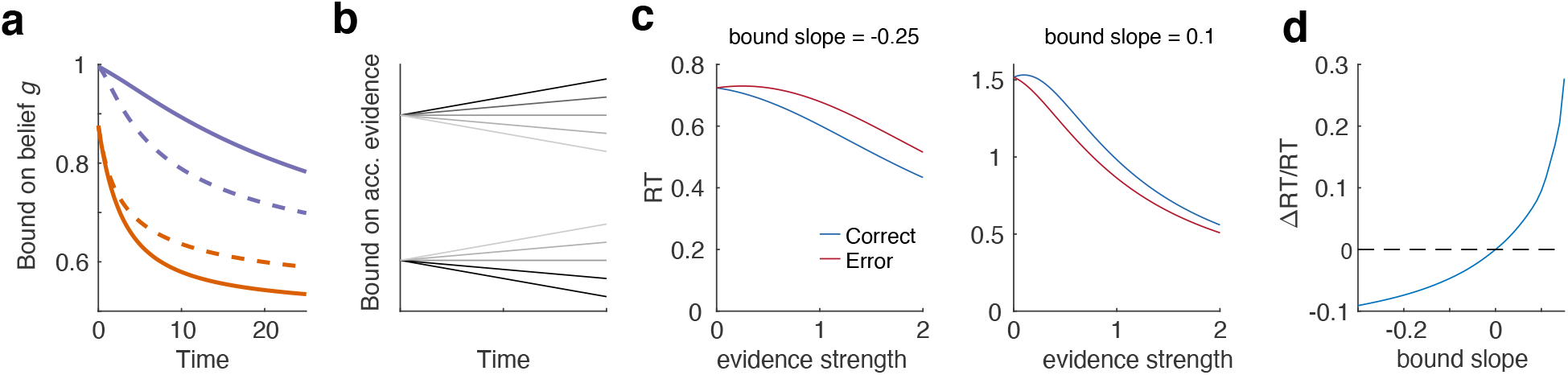
Relationship between the outcome-dependence of RTs and the slope of the decision bounds. **(a)** In solid lines, same decision bounds in belief as in Fig. 1e (excluding the lower value of the stakes). In dashed lines, belief bounds that would be obtained if the bounds on accumulated evidence were constant and equal to their value at t=0. For the orange case, the dashed curve is above the solid one, suggesting that the bounds on accumulated evidence are decreasing in time while, for the purple case, the dashed curve is below the solid one, suggesting that the bounds on accumulated evidence are increasing in time. **(b)** In order to understand the qualitative difference in the outcome-dependence of RT (∆*RT*/*RT* = (*RT*_*cor*_ − *RT*_*err*_)/*RT*_*total*_) in these two cases (Supplementary Fig. 2c), we considered a minimal model where the decision bounds in x are linear with variable slope and fixed intercept (i.e. same value at *t* = 0), such that the bounds can either be increasing or decreasing in time. **(c)** Two examples are shown for the mean RT of correct and error responses for different signs of the slope of the bound on accumulated evidence. Left: example with negative slope, for which the RT of errors is larger than the RT for corrects. Right: example with positive slope, for which the RT of corrects is larger than the RT of errors. **(d)** plotting ∆*RT*/*RT* as a function of the bound slope reveals that a change in the sign of the latter is sufficient to induce a change in the sign of the former, and that there is a one-to-one relationship between the two quantities. Interestingly, the relationship is highly non-linear.

**Supplementary Fig 5.**
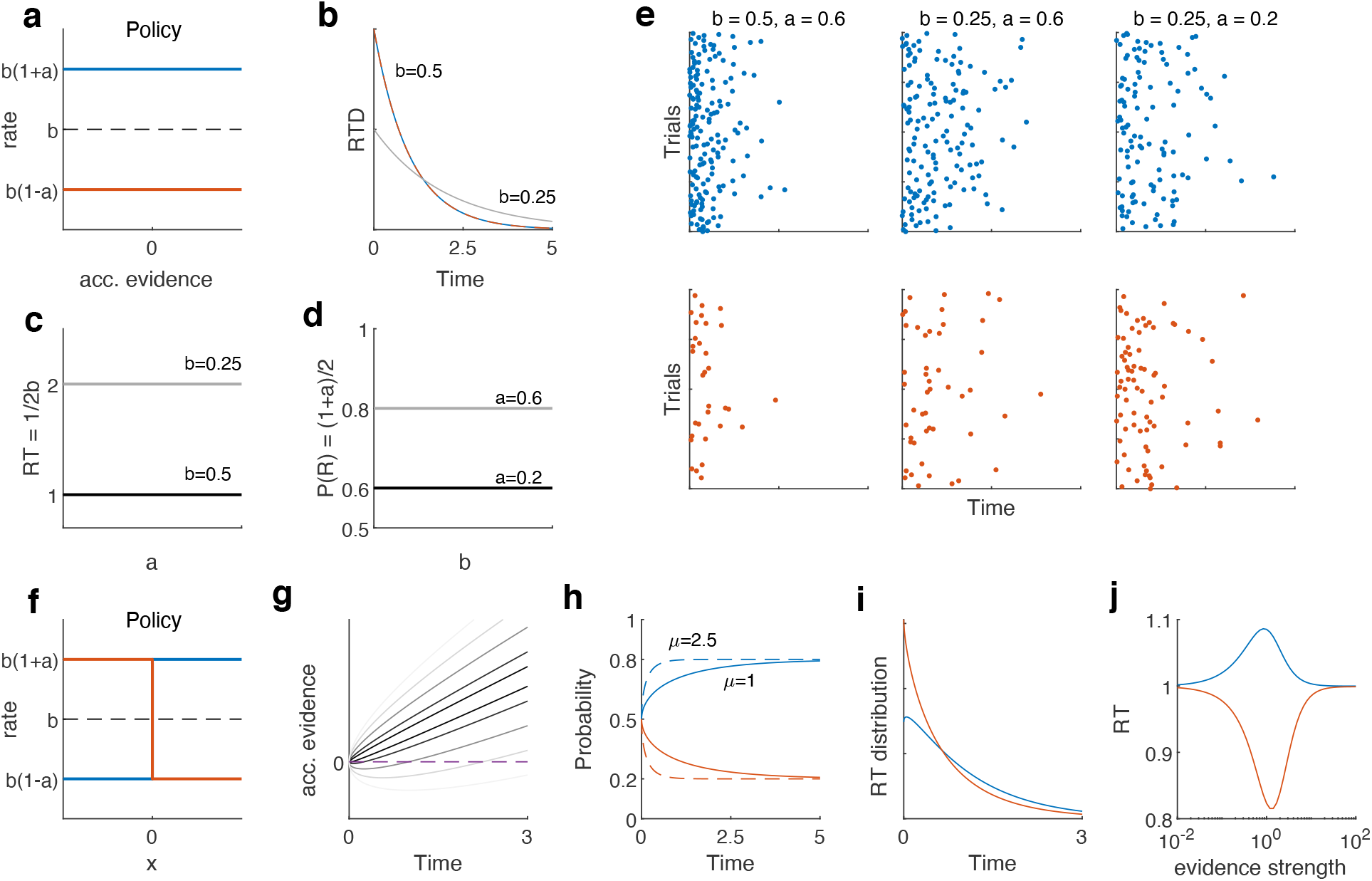
Properties of Poisson rate policies. **(a)** Example of a constant rate policy (homogenous Poisson race) as a function of the accumulated evidence, with the parameters *b* and *a* determining the sum and difference between the rightward (blue) and leftward (orange) rates. A choice is made towards the action whose policy commits first. **(b)** RTD produced by two constant rate policies with different values of *b*. The distributions are the same for rightward or leftward choices. **(c)** As a consequence, mean RTs only depend on *b* (*E*[*RT*] = 1/2*b*), and are constant as a function of *a*. **(d)** Conversely, the proportion of rightward choices only depends on *a* (*P*(*R*) = (1 + *a*)/2), and is constant as a function of *b*. **(e)** example trials visually demonstrating the results in (b-d). Top panels show rightward choices, bottom show leftward choices. In each column, the resulting RTDs are the same irrespective of choice. From the first to the second column, only *b* changes, so the choices overall become slower, but the proportion of rightward choices does not change. From the second to the third column, only *a* changes, so the RTDs stay the same but the proportion of rightward choices changes. **(f)** Example of a simple monotonic rate policy: the rates are step functions of the accumulated evidence, parametrized in the same way as before. As the accumulated evidence changes sign, the rightward rate transitions from low to high, and the leftward rate does the opposite. The sum of both rates is still constant and only depends on *b*. **(g)** Distribution of accumulated evidence as a function of time for evidence strength *μ* = 1. The lines represent quantiles of this distribution. **(h)** As the mass of the distribution of accumulated evidence shifts towards positive values with time, the conditional probability of a rightward choice increases and the conditional probability of a leftward choice decreases. Increasing *μ* (solid vs dotted lines) changes the speed of convergence towards the asymptotic values, which are still (1 ± *a*)/2. **(i)** This causes the RT distributions for rightward and leftward choices to be different, as they are the product of the probability of making a choice at a given time (which still only depends on *b* and it is the same as in (b)) times the conditional probability of a given outcome (h). In this example, *μ* = 1. **(j)** Mean RT conditioned on choice as a function of evidence strength (in log scale). The mean total RT is always the same because it only depends on *b*, but it differs conditioned on outcome as the evidence strength controls the shape of the curves in (h). Throughout panels (h-j), *a* = 0.6 and *b* = 0.5.

**Supplementary Table 1.**
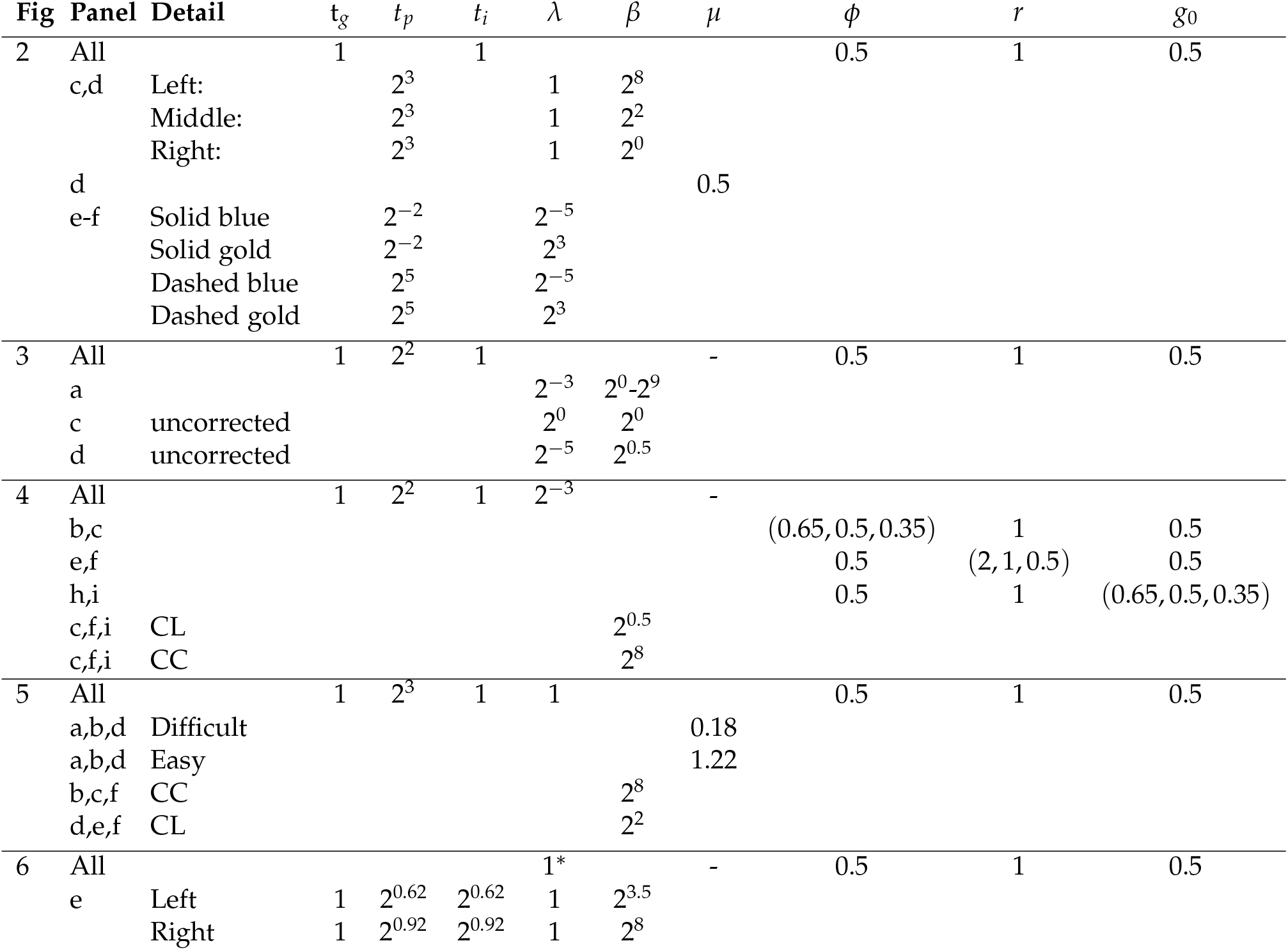
Parameters values for all figures in the main text. *In absolute units, not relative to *t*_*g*_.

| Fig | Panel | Detail | $t_g$ | $t_p$ | $t_i$ | $\lambda$ | $\beta$ | $\mu$ | $\phi$ | $r$ | $g_0$ |
| --- | --- | --- | --- | --- | --- | --- | --- | --- | --- | --- | --- |
| 2 | All |  | 1 |  | 1 |  |  |  | 0.5 | 1 | 0.5 |
| | c,d | Left: | | $2^3$ | | 1 | $2^8$ | | | | |
| | | Middle: | | $2^3$ | | 1 | $2^2$ | | | | |
| | | Right: | | $2^3$ | | 1 | $2^0$ | | | | |
|  | d |  |  |  |  |  |  | 0.5 |  |  |  |
| | e-f | Solid blue | | $2^{-2}$ | | $2^{-5}$ | | | | | |
| | | Solid gold | | $2^{-2}$ | | $2^3$ | | | | | |
| | | Dashed blue | | $2^5$ | | $2^{-5}$ | | | | | |
| | | Dashed gold | | $2^5$ | | $2^3$ | | | | | |
| 3 | All | | 1 | $2^2$ | 1 | | | - | 0.5 | 1 | 0.5 |
| | a | | | | | $2^{-3}$ | $2^0-2^9$ | | | | |
| | c | uncorrected | | | | $2^0$ | $2^0$ | | | | |
| | d | uncorrected | | | | $2^{-5}$ | $2^{0.5}$ | | | | |
| 4 | All | | 1 | $2^2$ | 1 | $2^{-3}$ | | - | | | |
|  | b,c |  |  |  |  |  |  |  | (0.65,0.5,0.35) | 1 | 0.5 |
|  | e,f |  |  |  |  |  |  |  | 0.5 | (2,1,0.5) | 0.5 |
|  | h,i |  |  |  |  |  |  |  | 0.5 | 1 | (0.65,0.5,0.35) |
| | c,f,i | CL | | | | | $2^{0.5}$ | | | | |
| | c,f,i | CC | | | | | $2^8$ | | | | |
| 5 | All | | 1 | $2^3$ | 1 | 1 | | | 0.5 | 1 | 0.5 |
|  | a,b,d | Difficult |  |  |  |  |  | 0.18 |  |  |  |
|  | a,b,d | Easy |  |  |  |  |  | 1.22 |  |  |  |
| | b,c,f | CC | | | | | $2^8$ | | | | |
| | d,e,f | CL | | | | | $2^2$ | | | | |
| 6 | All | | | | | $1^*$ | | - | 0.5 | 1 | 0.5 |
| | e | Left | 1 | $2^{0.62}$ | $2^{0.62}$ | 1 | $2^{3.5}$ | | | | |
| | | Right | 1 | $2^{0.92}$ | $2^{0.92}$ | 1 | $2^8$ | | | | |

## Supplementary Note

Here we present the mathematical derivation of our framework. To make the document self-contained, we include some theoretical background when needed.

### 1 Framework

#### 1.1 Markov Decision Processes: First exit formulation

We consider a Markov Decision Process^42,54,0^ (MDP) with a set of states *s* ∈ *S* and a set of admissible actions *a* ∈ *A*. The actions generate transitions between states according to a transition probability *T*(*s*^′^|*s, a*) which crucially is Markov in the states and actions. Associated to each transition, there is an immediate reward ℛ (*s, a*). The goal of the agent is to maximize the long term reward accumulated over the entire sequence of transitions. In order to do this, the agent is equipped with a control policy *u*(*a*|*s*), i.e., a decision rule to select actions in each state. The problem is then to find the optimal policy that ensures the largest possible long term reward.

In the “first-exit” formulation of the problem^42,54^, there is a set *S*_*T*_ ∈ *S* of “terminal” states which, once reached, terminate the process. The accumulated reward starting from state *s* and acting optimally thereafter, called the value function *Y*(*s*), can be written as

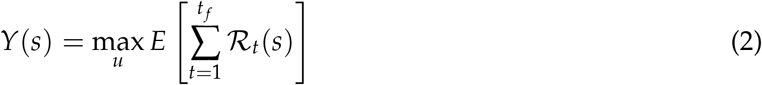

where ℛ_*t*_(*s*) is the reward obtained at time *t* having started from state *s* and following the policy, and the expectation is taken over any stochasticity in the process. *t* _*f*_ denotes the time step at which the first terminal state is reached (which is also a random variable). The value function follows a recursive relationship called the Bellman Equation^54,110^ (BE), which is given by

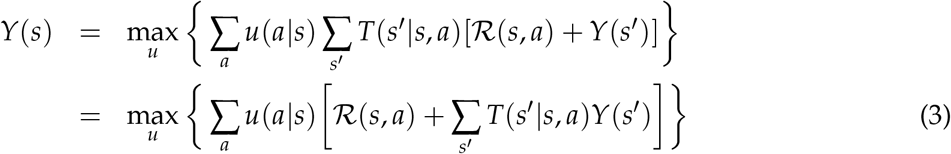

It can be shown that if the immediate cost ℛ (*s, a*) does not depend on *u*, the optimal policy must be deterministic^0^ and given by

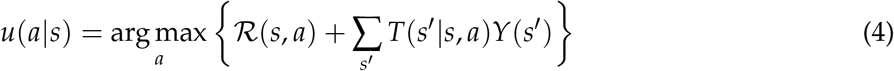

which allows rewriting the equation (3) as

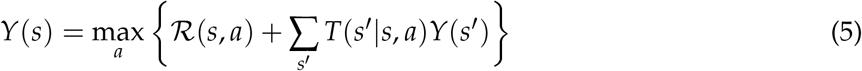

#### 1.2 Kullback-Leibler control

In the above formulation of the problem, standard in the field of Reinforcement Learning, there are no costs associated to the control of the agent. In the field of Optimal Control, two reasonable features are added: First, it is assumed that the agent is capable of behaving in the absence of control. Second, it is assumed that control is costly, and that this cost should be included in the optimization process. Formalizing this idea means considering *two* policies. One is a control-free default (or passive) policy *p*(*a*|*s*), which describes the behavior of the agent in the absence of control and which, in principle, bears no particular relationship to the goals of the agent. This passive policy is part of the specification of the problem and plays a role similar to that of the prior in statistical inference. The other one is the target optimal policy *u*(*a*|*s*) which describes the trade-off between long-term reward maximization and control costs.

The framework of KL control^53–55,109,0^ assumes a specific form for the cost of control, which is added to the immediate reward ℛ (*s, a*). The immediate consequence of choosing action *a* in state *s* under the optimal policy *u*(*a*|*s*) for an agent with default policy *p*(*a*|*s*) thus becomes

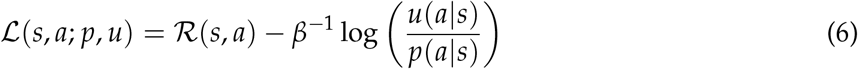

The constant *β*^−1^ measures the magnitude of the cost of control relative to the immediate reward, and can be considered a property of the agent. As we will see, when the cost of control is negligible (i.e., *β* → ∞), the attainment of long term reward dominates the behavior of the agent, and the optimal policy becomes identical to the one obtained in the standard MDP framework. However, when control is costly, the optimal behavior of the agent will tend to be constrained by its passive policy. Replacing ℛ (*s, a*) by ℒ (*s, a*; *p, u*) in the BE (5) one obtains

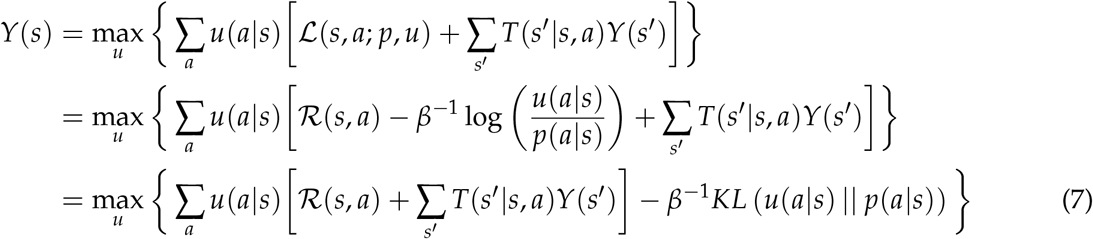

where *KL*(*p*||*q*) is the Kullback-Leibler divergence^56^ between distributions *p*(*x*) and *q*(*x*), which gives its name to this framework. Thus, the effective consequence of the control cost is to penalize the optimal policy by a quantity proportional to the dissimilarity between the two policies, as reasonably expected. The usefulness of this formulation can be appreciated when applying some further algebra on the above equation:

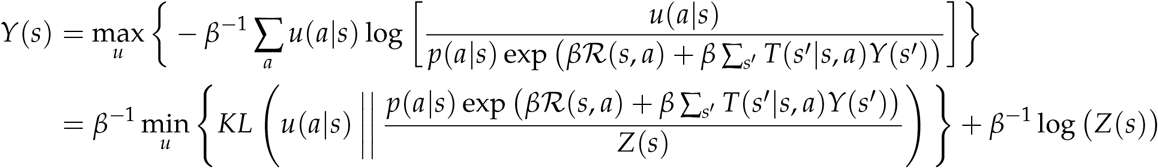

where the “partition function” *Z*(*s*) is defined as

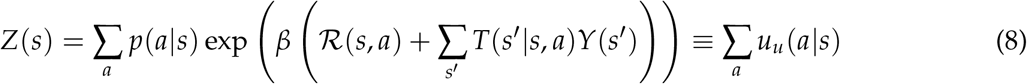

The control law results trivially from the minimization of the *KL* divergence, which is attained when its two arguments are equal, resulting in *KL* = 0. In this case, the BE reduces to

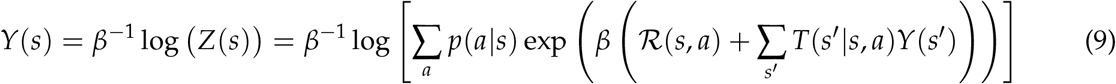

which is a self-consistency equation that specifies the value function. At the same time, the optimal policy is given by (from the condition *KL* = 0)

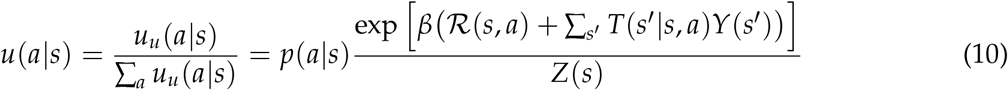

The last two equations (the equivalent of equations (5) and (4) above for the standard MDP) provide the solution to the KL control problem. It entails two important simplifications: First, unlike Eq. (5), Eq. (9) does not have a max_*a*_ operator and it is thus continuous. Second, once the value function in Eq. (9) is known, Eq. (10) provides an analytical expression for the optimal policy. This expression, furthermore, has an intuitive interpretation. Except for the normalization constant *Z*(*s*), the (un-normalized) optimal probability of choosing each action under the optimal policy *u*_*u*_(*a*|*s*) is proportional to the same probability under the passive policy *p*(*a*|*s*), with a proportionality constant that grows exponentially the net “action value” of the pair (*a, s*). One important consequence of this fact is that actions that have zero probability of being chosen under the default policy remain forbidden for the optimal agent. Thus, in KL control, control can only bias the probability of actions that were possible by default, it cannot create new actions *de novo*. This limitation can be seen as a price to pay for the mathematical simplification associated to the choice of the KL divergence as a measure of control cost.

#### 1.3 Equivalence between MDP and KL formulations

Equations (9) and (10) are instances of the functions LogSumExp and SoftMax respectively

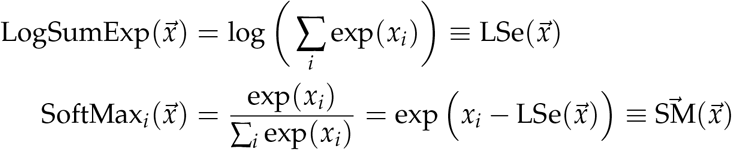

These are smooth versions of the *max* and *argmax* operators correspondingly. Notice that knowing the value of the first one facilitates the calculation of the second one, as it is the logarithm of the denominator. Expanding *Z*(*s*) in both (10) and (9)

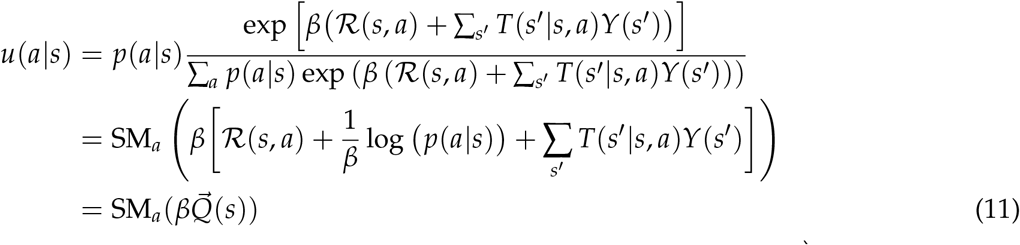

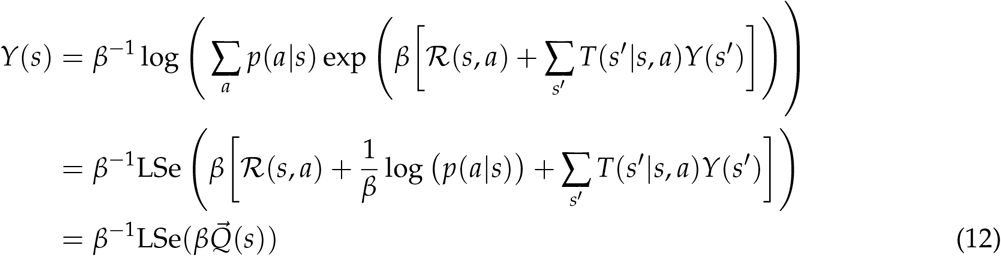

where we have defined the vector 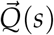 with components

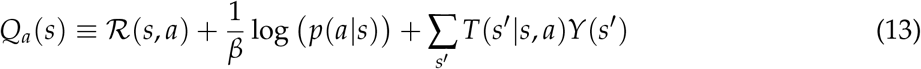

Each *Q*_*a*_(*s*) measures the *action value* of being in state *s* and performing action *a* including a measure of control given by the surprisal log(*p*(*a*|*s*)). Since this is an immediate cost, it is useful to include it in the immediate reward, which we thus redefine to be

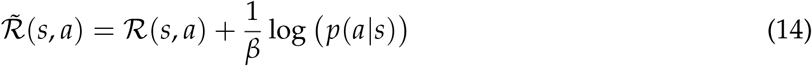

The second term is always negative, i.e., a cost. Thus, the more unlikely an action is in a particular state under the default policy, the lower the effective immediate reward in that state. This can be construed as a form of directed exploration towards the default. Using this modified effective immediate reward leads to the following natural expression for net action value *Q*_*a*_(*s*)

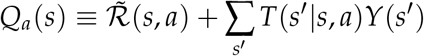

It’s straightforward to show that in the limit where *β* goes to infinity 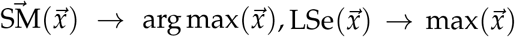, and 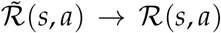, so that the value function and the policy converge to the MDP deterministic solution in Eqs. (4-5).

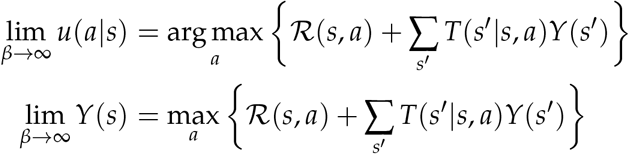

#### 1.4 Average-adjusted reward formulation

The first-exit formulation describes well a single trial in a decision making experiment, starting with stimulus onset and ending when the subject commits. However, a behavioral session contains many such trials, and requires a longer horizon. The standard way to deal with long (infinite) horizons is to use discounted rewards^0^. However, if the MDP is ergodic, there exists another approach which is well suited for describing a multi-trial decision-making experiment: reward rate maximization^42,43^. We explain this framework for a standard MDP, and later on show that it is trivially generalized to the KL control framework.

Let the MDP be represented by a discrete time, ergodic Markov chain^42^. We focus on the case where trajectories along the chain can be broken down in “trials”, with the beginning of each trial marked by the moment in which a certain initial state *S*_*I*_ is visited. *S*_*I*_ should be understood as a reference state, and the ergodicity of the chain allows us to make this choice arbitrarily.

Let us define

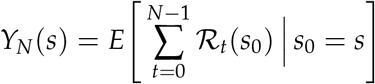

with the same interpretation as in (2) except that *N* is a fixed and sufficiently large natural number, and where we have omitted the explicit dependence of the sequence of rewards on the policy (we in general assume the optimal policy). Since the state *S*_*I*_ is always revisited, we can rewrite the equation separating it in two terms: one that captures the accumulated reward starting from state *s* until *S*_*I*_ is reached for the first time, and another one capturing the remainder of the accumulated value since the first time *S*_*I*_ is visited

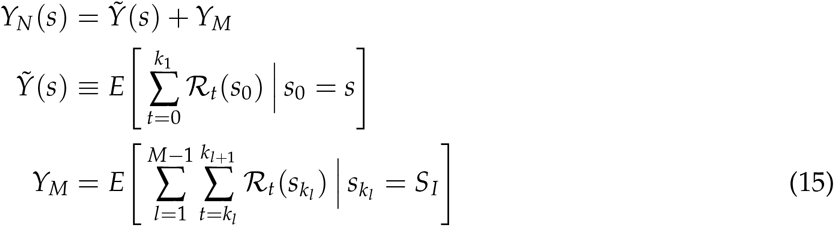

where *k*_*l*_ corresponds to the successive times that the state *S*_*I*_ is revisited, and the index *l* goes over trials, which go up to *M*. The conditioning on 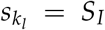 makes explicit that every trial is initiated at the state *S*_*I*_. Only the first term depends on the particular initial state *s* at time *t* = 0, while the second term is independent of *s* and linear in *M*. This is equivalent to separating the initial (in)finite horizon problem into an initial first-exit problem, and a reduced (in)finite horizon problem that consists of a collection of identical first exit problems.

The reward rate *ρ* is defined as

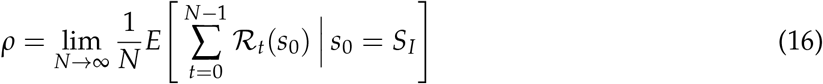

Since the chain is ergodic, the reward rate for any policy is constant and independent of the initial state, so we can choose *s* = *S*_*I*_. We can employ the same sort of separation of *N* into trials that we did above. Multiplying and diving by *M* in both numerator and denominator we obtain

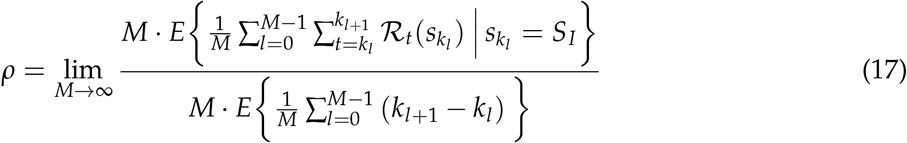

The expectation in the numerator corresponds to the average cumulative reward in one trial, while the expectation in the denominator corresponds to the average duration of a trial. Let us call these ⟨R⟩ and ⟨*T*⟩ respectively. Then we can simply write

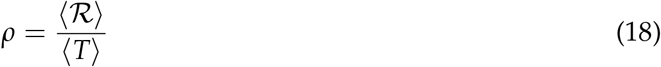

The reward rate *ρ* is also defined as the gain of the policy, since it measures the slope of the asymptotic linear dependence of total reward with the duration of the trajectory. Policies that obtain differences in total future reward which are constant as *N* grows will all have the same *ρ*, but it would be desirable to know which policies maximize these finite contributions as well. The relative value, or bias, *V*(*s*) of the policy, defined as

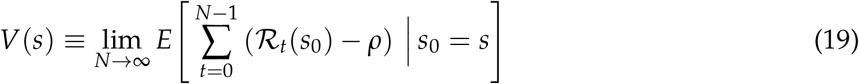

measures exactly this finite contribution, and is thus dependent on the initial state *s*. It can be shown^42,43^ that a policy that maximizes the relative value *V*(*s*) also automatically maximizes the reward rate *ρ*. In a multi-trial decision problem, *V*(*s*) can be written as

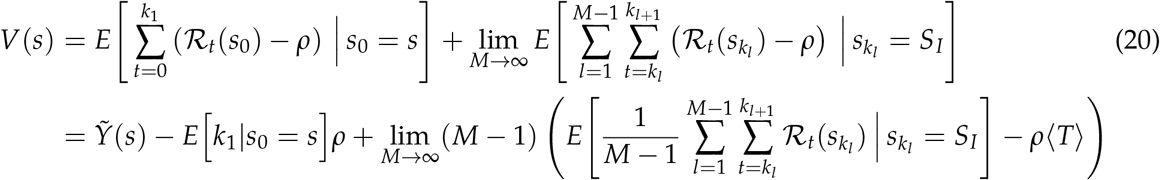

The expectation in the third term corresponds to ⟨ℛ ⟩ as we saw above, so that

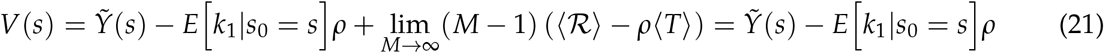

where the last equality uses the definition of *ρ* in Eq. (18). We now write the BE for the value function, taking the limit of *N* to infinity

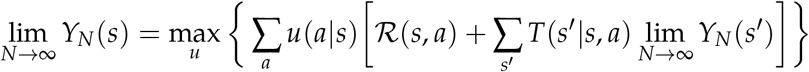

which, recalling that 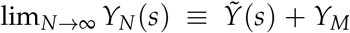, and that *Y*_*M*_ is a constant independent of *s*, becomes

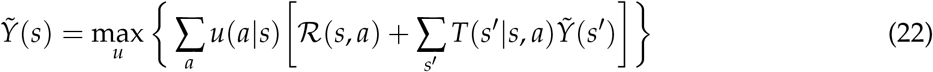

Using Eq. (21), this can be turned into a BE for the relative value *V*(*s*)

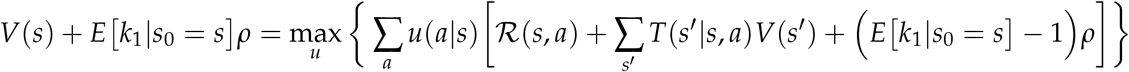

The last term uses the fact that the expected first exit time following a transition is exactly one unit less than the expected time before the transition, i.e.,

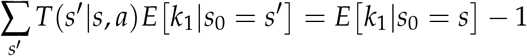

Since the expectation in this equation is already over trajectories obtained using the optimal policy, it can be taken outside the max operator, which leads to

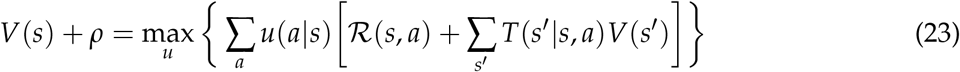

We have thus transformed an infinite horizon problem on *Y* into a first exit problem on *V*, with the additional constant *ρ* in the left hand side. Although *V*(*s*) is a relative value, we will refer to it as the value function from now on. The value of *ρ* is unknown, but we know it has to satisfy Equation (21). This implies that *V*(*S*_*I*_) needs to satisfy the following condition

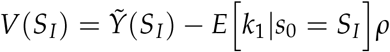

since *S*_*I*_ marks the start of a trial, the first term is the expected reward during one trial, ⟨R⟩, and the expectation in the second term is equivalent to the average duration of one trial, ⟨*T*⟩. Thus

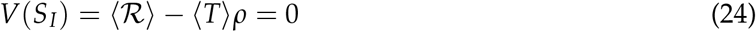

In practice, this equation is used to obtain the value of *ρ* self-consistently: We solve the BE (23) for a fixed value of *ρ*, use it to evaluate *V*(*S*_*I*_), and repeat this procedure adjusting *ρ* iteratively until *V*(*S*_*I*_) = 0. The value of *ρ* for which this condition is met is the average reward rate of the policy.

Notice that, mathematically, the BE (23) is identical to the BE for a first-exit problem with a “cost of time” *ρ*, i.e., a problem with an immediate reward for each (*s, a*) equal to ℛ (*s, a*) − *ρ*. Such BE is well defined for any arbitrary *ρ*. It is only if one wants to interpret *V*(*s*) as the relative value of an infinite horizon problem with asymptotic reward rate *ρ* that Eq. (24) needs to be satisfied.

Finally, because no assumptions have been made on the form of the immediate reward, the average-adjusted reward formulation for KL control is trivially obtained replacing ℛ (*s, a*) in the previous equations by Equation (6). The relevant expressions become

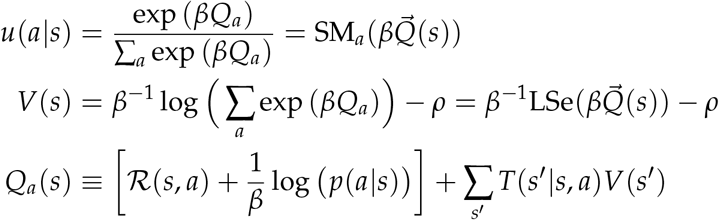

#### 1.5 Partial Observability: POMDPs

So far, we have been assuming that the agent has perfect knowledge about the states of the environment. In perceptual decision-making tasks, however, the challenge faced by the agent is precisely that the relevant states of the environment from the point of view of reinforcement are not directly observable, and have to be inferred through inference, based on stochastic observations. The appropriate mathematical framework to describe these situations is that of partially-observable Markov decision processes^16,111^ (POMDPs).

Qualitatively, this introduces the need for “information seeking” actions. In the problem we describe in the text, this action corresponds to the postponement of commitment, or waiting (which we have denoted by *W*). Although accumulation of evidence across time would seem to require memory and thus violation of the Markov assumption, it can be shown that the probabilistic belief of the agent (i.e., the posterior probability of the states given the full history of observations) can be updated recursively in a Markovian fashion, i.e., the agent’s belief in the current time-step is only a function of the current observation and the belief in the previous time-step^16,112^. Thus, formally, a POMDP can be construed as an MDP where states are replaced by beliefs^16^. If one uses the notation 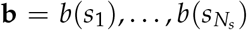 to refer to the relevant beliefs of the agent over the states *s*, then the POMDP formulation of our problem is very similar to the one we have previously described

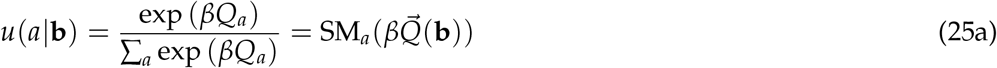

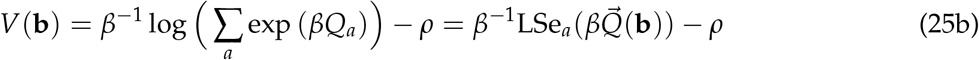

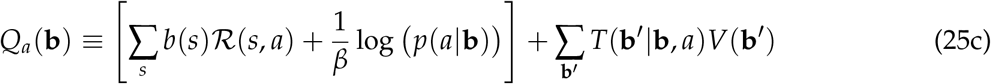

In this case, the immediate rewards ℛ (*s, a*) now correspond to the expected reward by the agent given its current beliefs. The quantity *T*(**b**^′^|**b**, *a*) replaces the transition probability *T*(*s*^′^|*s, a*) and describes the dynamics of belief induced by the dynamics of the agent-environment. Writing it in this form requires a marginalization over the observations that might be received if the agent performs action *a* with beliefs **b**, which will determine its subsequent belief state **b**^′^, as we now describe.

### 2 Modeling a binary choice over a continuous latent state

#### 2.1 States and transitions

The states in the task follow a continuous time Markov Chain (CTMC) as depicted in Fig. 1. The most important state is the “stimulus” state. This state is the only one that admits actions from the agent, and therefore the only one that offers immediate rewards ℛ (*s, a*). The possible actions are: waiting (*W*), choosing right (*R*) or choosing left (*L*).

The stimulus is characterized by a latent, continuous, unobserved feature (*μ*), which needs to be categorized by the agent as positive or negative. We define the task contingency to require *R* (*L*) when *μ* > 0 (*μ* < 0). We thus refer to *μ* > 0 (*μ* < 0) as the “right stimulus” – RS (“left stimulus” LS). If the agent waits, the current state is maintained (including the value of *μ*). But if the agent chooses *R* or *L*, the stimulus will end, and the task moves to a different state. In this transition, the agent will receive the ℛ (*s, a*) which depends on whether the choice is correct or an error. Let us define the reward for being correct as *R*_*w*_ and the reward for being incorrect as 0 (later we will show that the framework is invariant under the scaling of this reward). In addition, the following state will be different depending on a correct or an incorrect response. After a correct choice, the task moves directly to the “Inter Trial Interval” (ITI), with duration *t*_*i*_. After an error, the task advances to a “Time Penalty” state (TP), with duration *t*_*p*_, after which the task proceeds to the ITI. The time penalty becomes another incentive for the agent to choose correctly – besides the potential reward as the goal is to maximize the reward rate.

We can construct a table to make the different state transitions and payoffs from the Stimulus state more explicit.

State transitions (*T*(*s*′|*s, a*)):

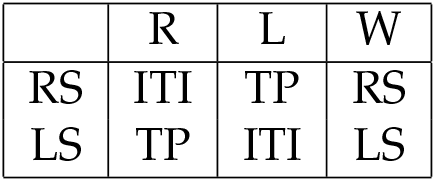

Payoff matrix (ℛ(*s, a*)

Payoff matrix ( $\mathcal{R}(s, a)$ ):
|  | R | L | W |
| --- | --- | --- | --- |
| RS | $R_w$ | 0 | 0 |
| LS | 0 | $R_w$ | 0 |

Notice that the model of transition probabilities defined in this way is *deterministic*, as each combination of states and actions always gives raise to the same successor state.

#### 2.2 Default policy

We model the agent’s default response tendencies through a constant, stimulus-independent probability of commitment per unit time

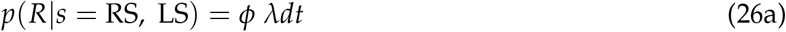

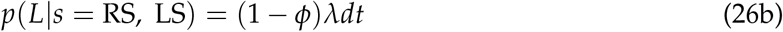

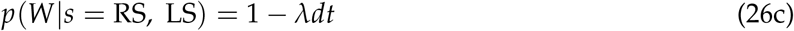

where *λ* is the rate of commitment in Hz. In most cases we assume responding is unbiased, i.e., *ϕ* = 1/2, although we relax this assumption when modeling sequential dependencies.

#### 2.3 Inference

In this section we derive expressions for the temporal evolution of the agent’s belief during the stimulus presentation. The relevant belief, *g*, is a scalar between 0 and 1 representing the probability that the stimulus is rightward, i.e., that *μ* > 0. Since the latent state *μ* is continuous, the agent calculates *g* indirectly by first estimating the posterior distribution of *μ* given the observations. If at time *t* the posterior over *μ* is *p*(*μ, t*) then

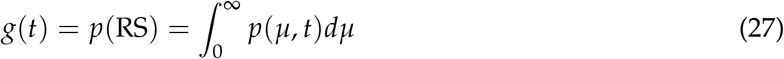

The stimulus emits a stream of temporally uncorrelated Gaussian observations. It is simpler to initially assume an observation is emitted every short interval ∆*t* with probability

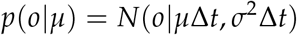

Without loss of generality, we also assume the prior distribution over *μ* is also Gaussian, and equal to

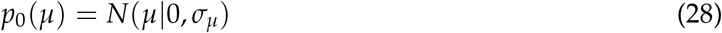

which implies an initially neutral belief

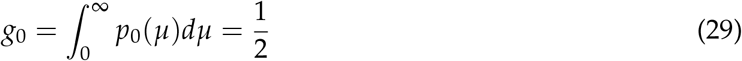

Because both the prior and the likelihood are Gaussian, the posterior is always Gaussian. It is easy to show that the posterior after *n* observations is equal to^18^

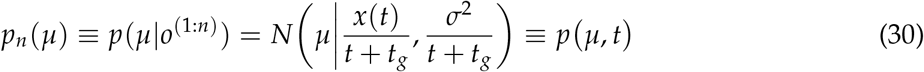

where

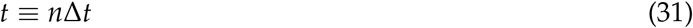

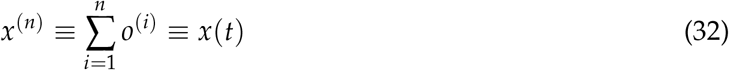

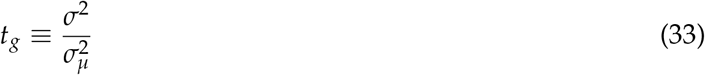

and thus that the belief at this time is

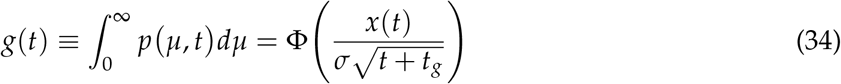

where Φ is the standard cumulative Gaussian distribution. These expressions show that the total accumulated evidence and elapsed time at a given moment are sufficient statistics to specify the posterior over the latent state, and thus also the agent’s belief. Note also that when *t* = *t*_*g*_ the posterior uncertainty is 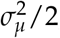, which is half of its initial value. The constant *t*_*g*_ thus measures the characteristic timescale of the inference process.

It is convenient to renormalize the variables of the problem to make them dimensionless as follows

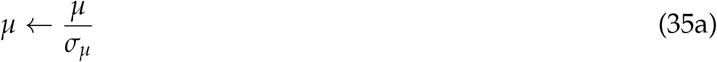

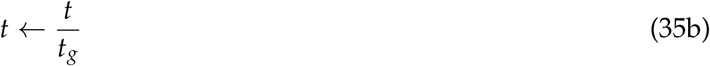

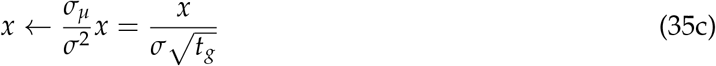

which simplifies the expressions of the prior, posterior and belief to

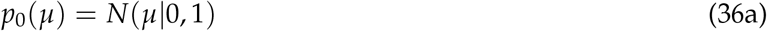

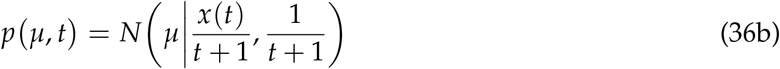

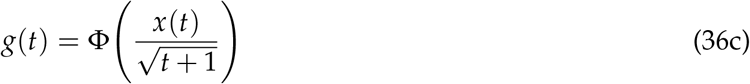

We also consider the problem in continuous time, by making the time interval infinitesimal, i.e., ∆*t* → *δt*, in which case the accumulated evidence *x*(*t*) becomes a continuous Markov process^113^ (CMP) with Langevin equation

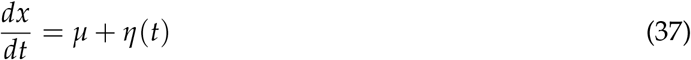

where *η*(*t*) is a white noise process. To proceed, we need to obtain the posterior predictive distribution of *x*(*t* + *δt*)^114^, meaning *x* in the following infinitesimal step, given *x*(*t*). If *μ* was known, this would be simply:

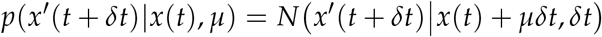

However, the agent does not know *μ*, only its probability distribution *p μ, t*. Marginalizing over *μ* one obtains

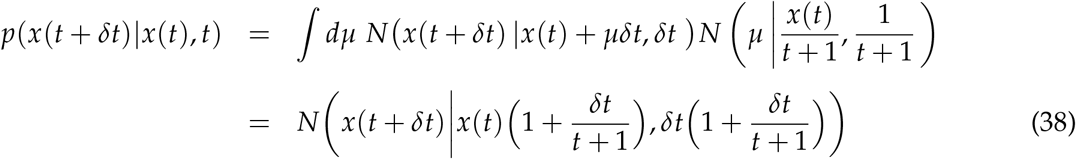

Thus, the prediction of *x*(*t*) follows another CMP with equation

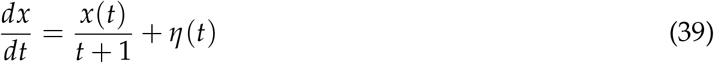

To obtain the corresponding (predictive) dynamics of belief, we write the last expression as a stochastic differential equation

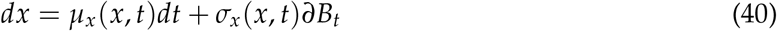

with

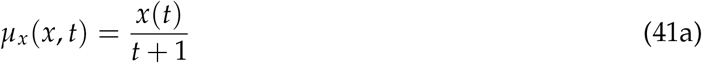

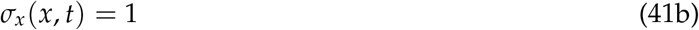

and invoke Ito’s lemma^115^, using the fact that *g*(*t*) is a monotonic function of *x*(*t*) (Eq. 36c)

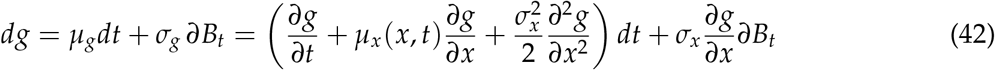

After some algebra, one obtains that the terms inside the parenthesis of the RHS of Eq. 42 cancel each other so *μ*_*g*_ = 0. Thus, the predictive distribution of the belief increment is normal and given by

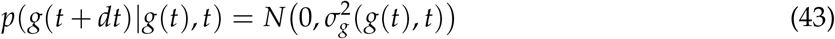

with

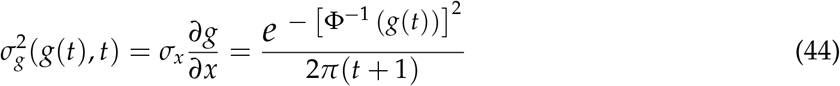

#### 2.4 Dimensional analysis

As just shown, it is natural to measure elapsed time since stimulus onset in units of *t*_*g*_. Under an appropriate renormalization (Eq. 35), inference then becomes parameter-free. The remaining free parameters of the problem are *λ, β, t*_*i*_, *t*_*p*_ and *R*_*w*_. Constants with units of time or rate should be measured in units of *t*_*g*_

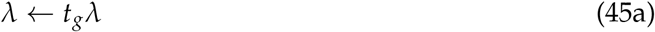

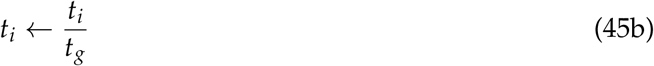

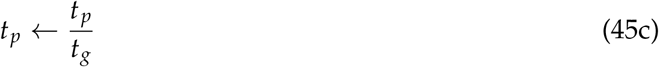

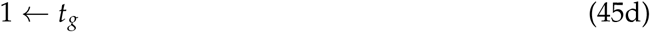

Thus, *λ*, for instance, represents the expected number of default choices in the time interval *t*_*g*_ necessary to roughly estimate the stimulus, on average across difficulties. The control limitation *β*^−1^, has units of reward, and so has the value function (Eq. 7). It is natural, thus, to use

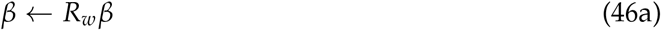

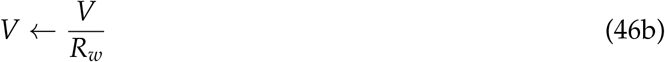

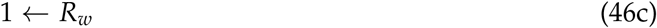

Conceptually, the fact that *β*^−1^ has units of reward means that control limitations are to be measured relative to the motivation of the agent (see text).

Although the reward rate *ρ* is not a free parameter, the preceding arguments imply that the mathematical framework is invariant with respect to changes in the magnitude of *t*_*g*_ and *R*_*w*_ if one also defines

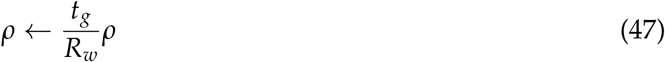

This scaling of reward rate is consistent with the fact that making the task more difficult (increasing *t*_*g*_) leads to lower reward rate, and vice versa (see text). Thus, the reward rate, but not other aspects of the policy, depends on *t*_*g*_ itself beyond its effect of rescaling the other free parameters of the model.

From now on, we always work with dimensionless parameters.

#### 2.5 Bellman Equation

In order to apply the average-adjusted reward rate framework, one needs to select one state as a reference, preferably one that is fully observable. In a sensory discrimination task, the natural state is the onset of the stimulus, which then has by definition a value of zero (Section 1.4). By backward induction, one can then obtain the value of the ITI and TP states. Since in these states there are no payoffs and the actions do not trigger any transitions, the value lost in those states is equal to the reward rate times their duration

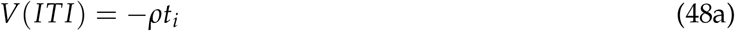

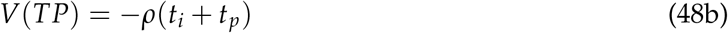

The agent transitions into these observable states from the stimulus state after committing to one of the two possible responses, with probabilities

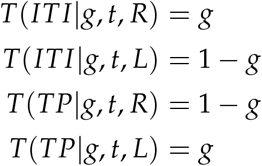

If the agent does not commit, it will remain in the stimulus state and its belief will evolve from the expected stimulus observations as described in the last section

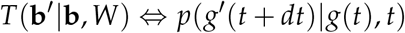

One then needs to compute the infinitesimal evolution of the expected value of the stimulus state. Because the predictive belief distribution (Eq. 43) depends explicitly on both *g* and *t*, the value of the stimulus state is also a function of belief and time since stimulus onset. The required expression is

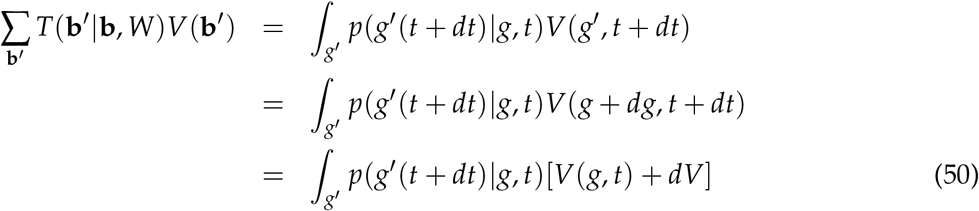

The value differential can be computed using Ito’s lemma again:

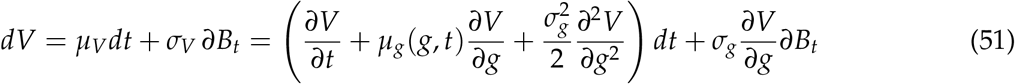

Since we are just looking for the expectation, we need to concern ourselves only with the term *μ*_*V*_, and since *μ*_*g*_ = 0, we obtain

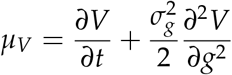

which implies

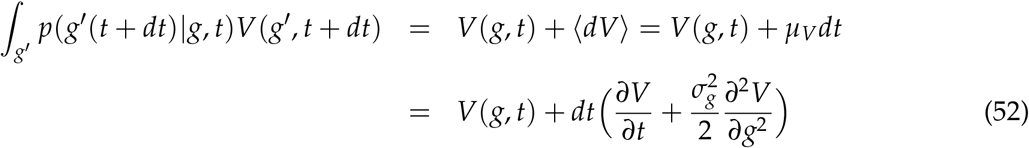

These expressions permit an evaluation of the actions values *Q*_*a*_, which in fact always appear exponentiated in the Bellman Eq. From Eq. 25c we can write

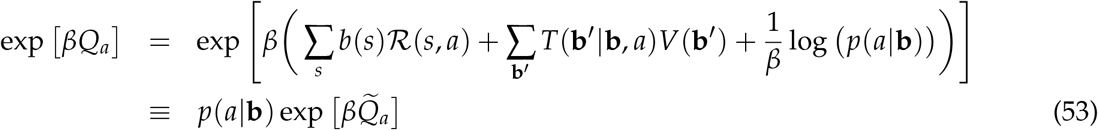

where

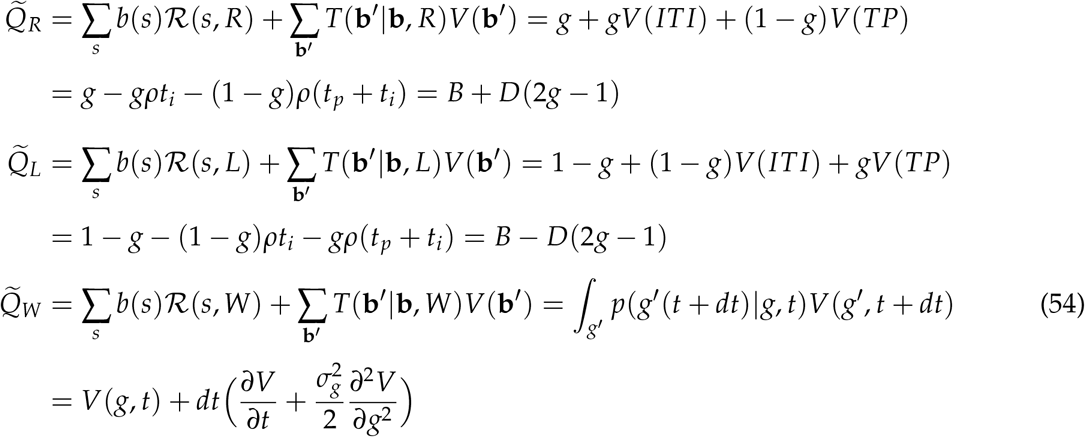

and where we have defined

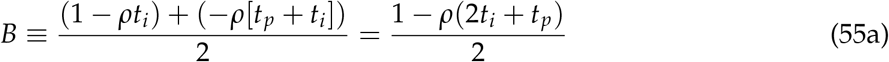

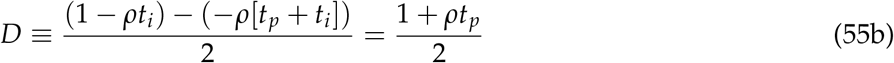

The quantity *B* represents the average immediate value across the two choices, and *D* represents their “contrast”. We can now write an expression for the Bellman Eq. (25b) in continuous time

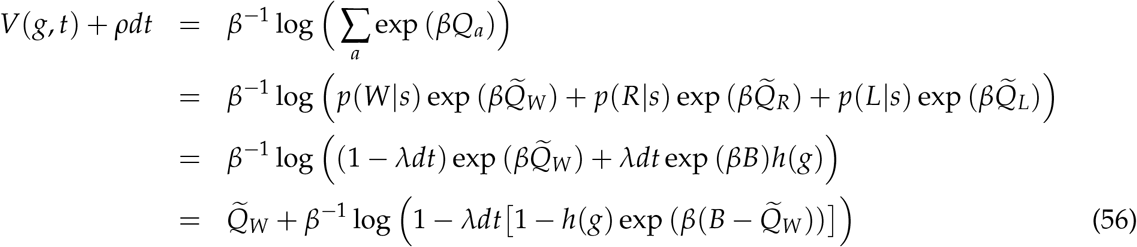

where we have defined

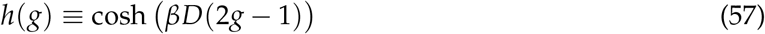

Substituting the expression for 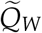 in Eq. 54, and developing to first order in *dt* we obtain

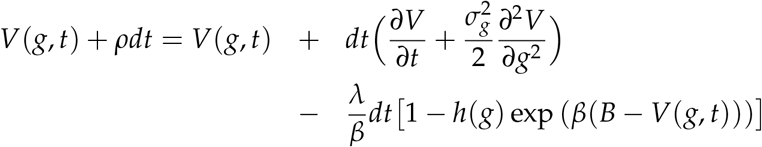

which leads to

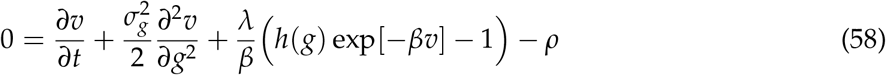

This is a second order, non-linear partial differential equation which *v*(*g, t*) ≡ *V*(*g, t*) − *B* (the value function during the stimulus presentation relative to the action-value *B* of a completely uncertain choice) must satisfy. To solve it, one needs to specify initial and boundary conditions.

##### 2.5.1 Initial and boundary conditions

The “initial” condition is in fact the value of the function at an unknown, large time *T*, because Eq. (58) needs to be solved backwards in time. This is the standard approach in Dynamic Programming, as the value of a state depends on the value of the options available in the future, not in the past. A first approximation can be obtained by considering the limit of Eq. (58) as *t* → ∞. In this limit, *v* should be stationary. It might seem counterintuitive that *v*, which represents the value of the stimulus state during which the agent is, by definition, uncommitted, can acquire a stationary value, given that the agent is paying a cost of time *ρ*. Intuitively, this is possible because after a sufficiently long time since stimulus onset, the probability that the agent will not have committed is vanishingly small. The value of being uncommitted at that state is thus given by the consequences of an immediate choice, which do not depend on the time elapsed since stimulus onset. Thus, one can impose ∂_*t*_*v* = 0. Since 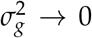 as *t* → ∞ (Eq. 44), the Bellman Eq. 58 in this limit becomes algebraic

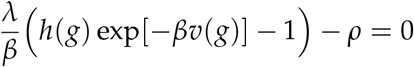

The steady-state value, which we denote by 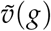 is thus given by

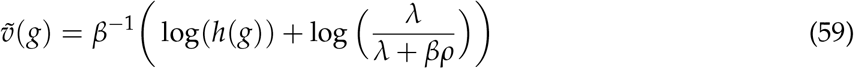

Qualitatively, the first term in this Eq. represents the value associated to an immediate choice, and the second term, which is negative, represents the loss in value – driven by the control limitations of the agent – derived from not being able to instantaneously choose the most valuable option, a delay that is always costly as long as *ρ* is non-zero. As such, this control cost vanishes when *λ* → ∞ (because then the agent spontaneously chooses without delay) or when *β* → ∞ (because in this case there are no control limitations).

In principle, one can initialize the value function at a sufficiently long time *T* as 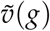 and numerically iterate backwards (see below). But this procedure is inefficient because the value of *T* necessary for 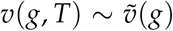 is very large. One can make this process more efficient by finding the first perturbative correction to 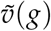. We seek a singular perturbative expansion of the form

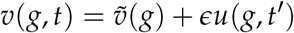

where *t*^′^ ∼ *O*(1) and *ϵ* ≡ *t*^′^/*t*. The function *u*(*g, t*^′^) describes the asymptotic temporal evolution of the value function discounting for the change of scale necessary to focus on infinitely long times. Changing variables in the Bellman Eq. 58 from *t* to *t*^′^ and neglecting terms ∼ *O*(*ϵ*^2^) one obtains, after some algebra

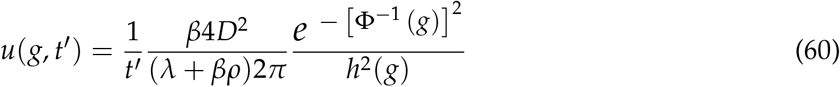

which leads to our desired result

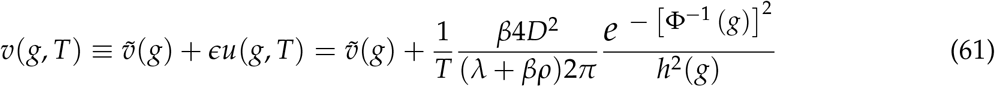

One can use this equation to provide an estimate for the long time *T* at which this asymptotic solution is expected to be accurate, by imposing that the overall scale of the second term is sufficiently small compared to the scale of the first one. Because *u* is maximal at *g* = 0.5 and the range of 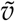 corresponds to 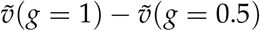, we impose that

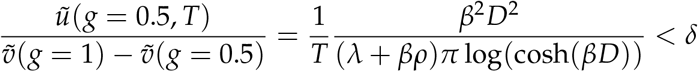

where *δ* ≪ 1 is a numerical tolerance (we typically choose *δ* = 10^−5^). Thus, we find

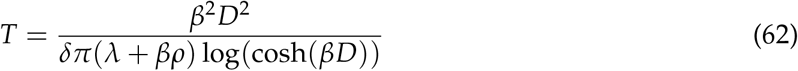

In practice, our numerical solution for the Bellman Eq. proceeds as follows. At first, given *δ*, we compute *T* using Eq. 62, and define *T*^′^ = 1.2 *T*. We then compute the asymptotic value function *v*(*g, T*^′^) using Eq. 61. In order for *T* and *T*^′^ to be adequate, we impose two conditions: (*i*) the numerically obtained value function *v*(*g, t*) for 0 < *t* < *T* should be the same when using *v*(*g, T*^′^) or *v*(*g*, 1.2 *T*^′^) as the asymptotic solution, and (*ii*) we update T to be equal to the numerically obtained (see below) 0.999 quantile of the reaction time distribution of the policy. These conditions guarantee that the asymptotic solution is truly asymptotic and does not affect any of the results, and that the numerical procedure is efficient in the sense of not calculating *v*(*g, t*) for values of time that the policy never visits. Usually, satisfying these solutions requires modifying iteratively the values of *T* and *T*^′^ a few times.

The boundaries correspond to the values *g* = 0 and *g* = 1, meaning full certainty that the stimulus state is *SR* or *SL* respectively. In these conditions, barring limitations in control, the agent should always choose immediately, as we argued in deriving 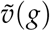. Since, as we showed in Eq. 59, the cost induced by these limitations is time independent, the value function in conditions of certainty is constant and equal to 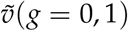

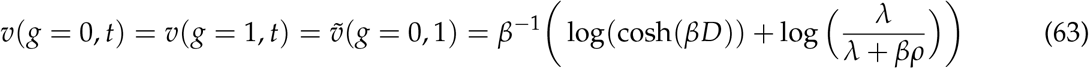

##### 2.5.2 Calculation of reward rate

As explained above, the reward rate *ρ* must be calculated by imposing that the relationship *V*_0_ ≡ *V*(*g*_0_, 0) = 0 be satisfied. We do this by an iterative procedure, starting with an initial guess and changing *ρ* repeatedly using the bisection method until the initial condition for the value function is satisfied. We find that *V*_0_ is monotonic in *ρ* and smooth, so the procedure is not problematic. Given that *ρ* has to satisfy

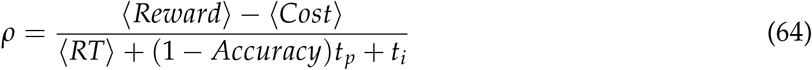

then, for the bisection method, we consider initial upper and lower bounds equal to

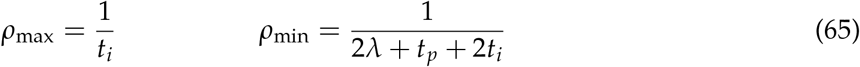

The expression for *ρ*_min_ is the reward rate of an agent following the default policy, which is a lower bound for any policy with *β* ≠ 0.

#### 2.6 Optimal Policy

Once the value function *v*(*g, t*) and *ρ* are known, it is possible to calculate the optimal policy from Eq. (25a)

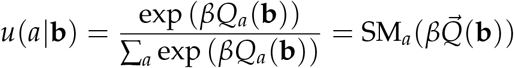

The denominator in this expression is obtained from the definition of the value function

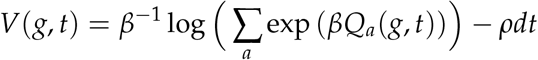

so that

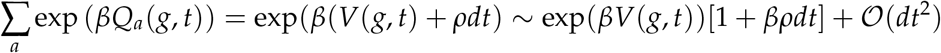

One can therefore write

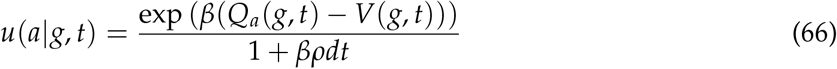

The quantity *A*_*a*_(*g, t*) = *Q*_*a*_(*g, t*) − *V*(*g, t*) in the exponent is known as the *advantage*^0,116^ of action *a*. It measures the excess or deficit in expected value from choosing an action relative to the average of all available actions following a given policy.

For the actions *a* = *R, L*, the exponential of the action-value is proportional to the instantaneous probability of commitment under the default policy (see, e.g., Eq. 56), and thus also proportional to *dt*. Using the expressions for *Q*_*R,L*_(*g, t*) in Eqs. 53-54, and developing to first order in *dt* one obtains

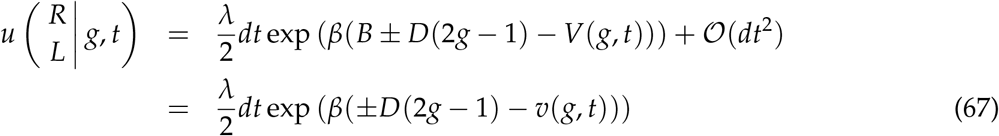

which corresponds to Eq. 4 and Fig. 2 in the text. The policy for the remaining action *W* is now obtained by normalization, i.e., *u*(*W*|*g, t*) = 1 − *u*(*R*|*g, t*) − *u*(*L*|*g, t*).

It’s informative to write the expression for the optimal policy for each choice as the product of the probability of commitment (which we denote as 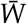), times the probability of choosing *R, L* given that a commitment has been made

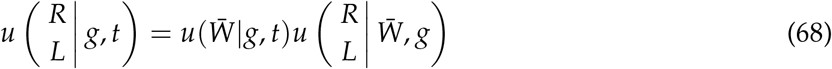

The probability of commitment is simply given by the sum of *u*(*R*|*g, t*) and *u*(*L*|*g, t*)

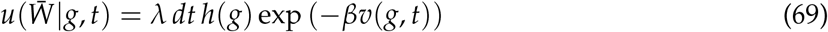

which leads to

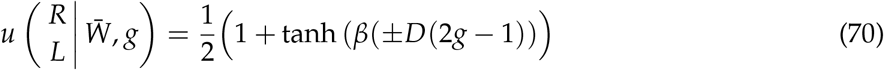

The complexity of the optimal policy is thus contained in the commitment decision, which depends on the time-dependent value of the stimulus state that requires solving the Bellman Eq. Once a commitment has been made, each action is selected through a simple, time-independent sigmoidal function of belief, which the steepness of the sigmoid an increasing function of the control ability *β* of the agent, and the “contrast” in value *D* between the two choices.

#### 2.7 Accuracy, reaction time and decision confidence

The optimal policy depends on belief and time, but the subject’s belief is unobservable. In each trial of an experiment, the subjects provide a binary choice, a reaction time and a measure of decision confidence. We seek a mathematical expression for the probability that, for a given stimulus difficulty *μ*, the subject will choose action *a* = *R, L* at time *t* = *RT* with a given level of confidence *g* = *DC*.

It turns out that it is simpler to work with the temporal evolution of accumulated evidence *x*(*t*) than with the subject’s instantaneous belief *g*(*t*), which does not represent a problem since there is a one-to-one monotonic relationship between them (Eq. 36c). It is important to note that, for this analysis, the dynamics of the accumulated evidence *x*(*t*) is given by Eq. 37 under a *known* stimulus *μ* (since *μ* is controlled by the experimenter), and not by the predictive distribution of *x*(*t*) induced by Eq. 39. The predictive distribution is needed to compute the optimal policy (which only assumes prior knowledge of the stimulus distribution *p*_0_(*μ*), Eq. 36a). Then, this policy is applied in all trials, under experiment-controlled sensory evidence characterized by *μ*.

We will slightly abuse notation and denote as *u*_*R,L*_(*x*(*g*), *t*)*dt* the optimal policy Eq. 67 as a function of *x*, and as 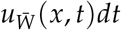 the corresponding probability of commitment. Because *x*(*t*) is a CMP, there is a Fokker-Plack Equation (FPE) associated with Eq. 37, which describes the dynamics of evidence accumulation in the absence of commitment

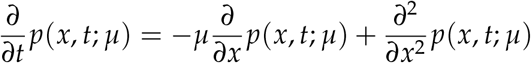

In the presence of a choice policy, the distribution 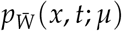, defined as the density at (*x, t*) given that the agent has not yet committed, will lose mass at a rate proportional to the rate of commitment 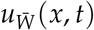 (Eq. 69), and will thus satisfy the following FPE

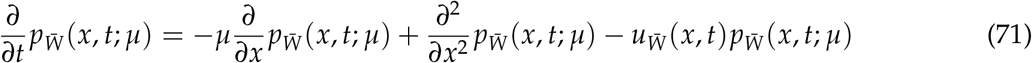

The probability mass that escapes is precisely what is needed in order to compute the probability density of committing to action *R, L* at time *t*, with accumulated evidence *x*.

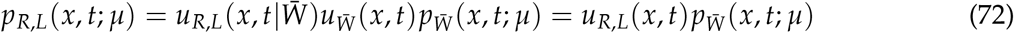

Eq. (71) needs to be solved under “natural” boundary conditions, since the domain is unbounded (see next section), and with initial condition

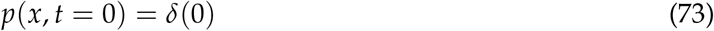

Given *p*_*R,L*_(*x, t*; *μ*), all quantities of interest can be computed through marginalization. For instance, the psychometric function, RT distributions and chronometric functions are given by

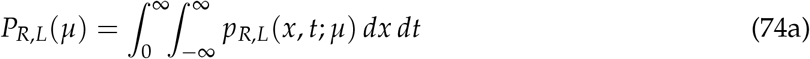

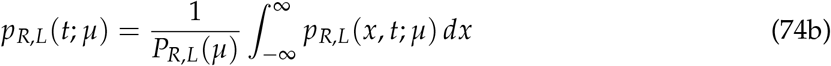

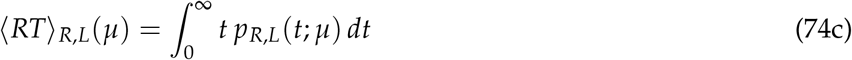

Similar expressions can be obtained for the distributions of decision confidence and their means.

##### 2.7.1 Decision lapses

We compute lapse rates by evaluating the value of 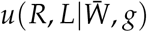 (Eq. 70) at *g* = 1 and *g* = 0 respectively. These are symmetrical for an unbiased default policy

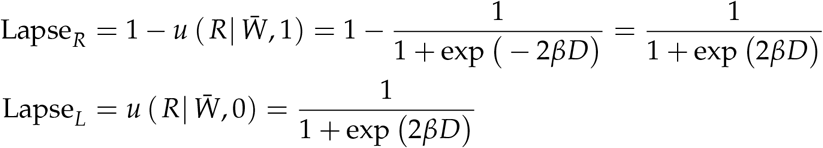

Intuitively, although computing the psychometric function in general requires the procedure described in the last section, lapses correspond to the limit *μ* → ±∞, and in this limit the situation simplifies. The distribution of *g*(*t*) effectively becomes a delta function at *g* = 0 or *g* = 1 instantaneously, and the rate is thus purely limited by the value of 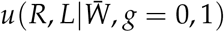.

### 3 Numerical methods

#### Value Equation

We solve Eq. 58 using finite differences, in particular the Method of Lines (MoL)^117,118^. Here, one uses algebraic approximations to spatial derivatives, and solves the resulting algebraic system of ordinary differential equations (ODEs) using standard packages.

We approximate

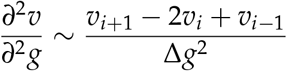

where *i* is an index that designates a position along a grid in *g* and ∆*g* is the spacing in *g* along the grid, which is assumed constant. We will consider that the grid as *M* points, so *i* = 1 and *i* = *M* are the extremes. These points will correspond to the boundary conditions, and we will need to treat them separately. With this discretization, the value equation takes the following form

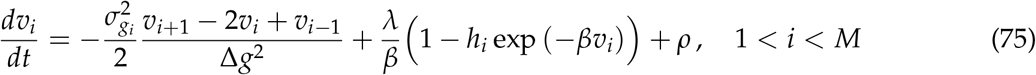

with

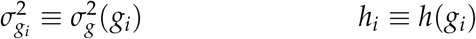

For the boundary conditions we have

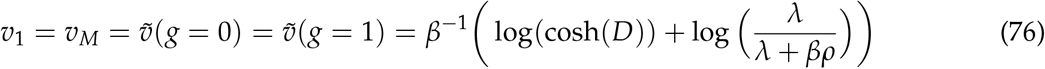

and the initual condition is just

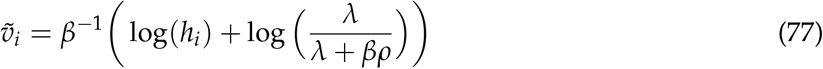

Eqs. 75 can be written in matrix form as

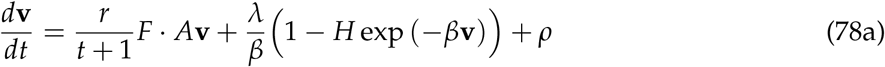

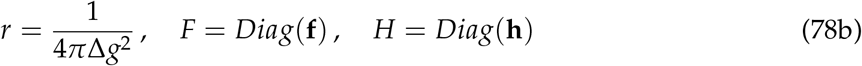

where

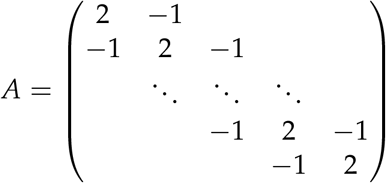

and where we have defined

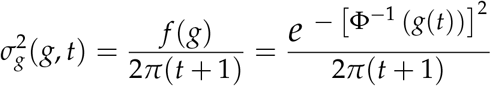

We typically use *M* = 501.

#### 3.2 Fokker-Planck Equation for Behavioral Predictions

Several techniques have been developed to maintain the positivity and the total probability of FPEs^119–121^. We use the Chang-Cooper method^119^, providing only enough details to keep our description self-contained. We start by writing the FPE 71 in flux form. The presence of the sink term does not allow achieving the canonical flux form, but nonetheless we can write

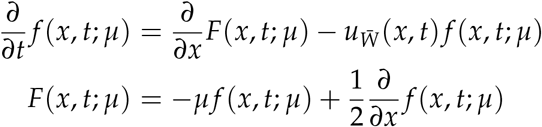

Where we have changed the notation 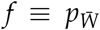, and *F* is the flux. Now we discretize *x* in a grid *x*_*i*_ such that ∆*x* = *x*_*i*+1_ − *x*_*i*_, with *x*_*i*±1/2_ ≡ *x*_*i*_ ± ∆*x*/2, and consider the discretization of the flux derivative

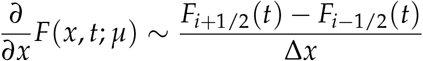

The goal is to choose the quantities *F*_*i*+1/2_(*t*) such that the positivity and normalization of the solution is guaranteed. This is achieved by setting

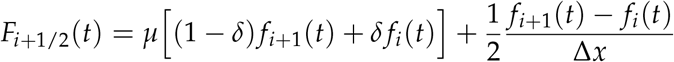

where we have defined *f*_*i*_(*t*) ≡ *f* (*x*_*i*_, *t*), and *δ* takes the value

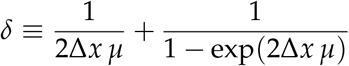

(see [120, 121] for details). This is valid for interior points. Regarding the boundary conditions, the physical domain of *x* is unbounded, but we have to limit it somehow in order to be able to compute the solution. The approach that we will use is to apply an absorbing barrier at ±*x*_*B*_, that is permeable to the sink force but captures the probability mass that diffuses into it. *x*_*B*_ > 0 is sufficiently large as to not disrupt the solution inside the barrier significantly^122^, given that probability mass is escaping due to the sink term. This is implemented reducing the equation at the boundaries *i* = 0, *i* = *M* to the following form

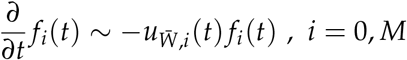

The resulting algebraic equation in vector-matrix form is

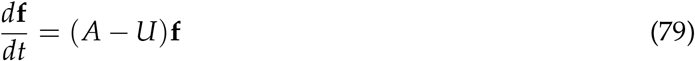

with *A* equal to

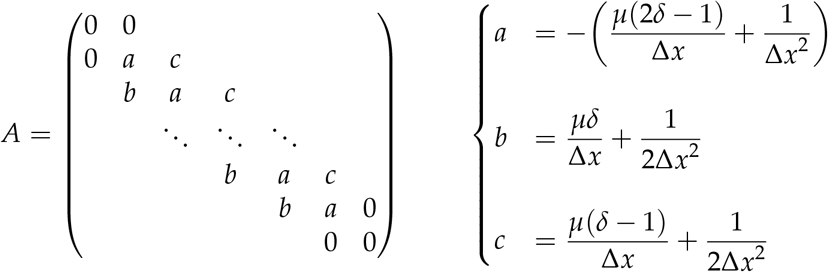

and 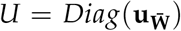. As for the initial condition, it is somewhat problematic to numerically implement a Dirac delta. A practical approach is to approximate the sink term as constant for a brief time interval ∆*t*, so then the PDE has an analytical solution:

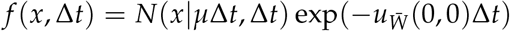

We typically use ∆*x* = 0.01 and ∆*t* = 0.0002.

##### 3.2.1 Determining *x*_*B*_

The limits *x*_*B*_ should be chosen such that the impact of using finite boundaries is negligible on the solution of the FPE. We choose *x*_*B*_ as to limit the maximum belief achieved. Given a certain belief cutoff defined as 1 − *δg*, with *δg* ≪ 1 (typically *δg* = 10^−3^ will be reasonable), we have

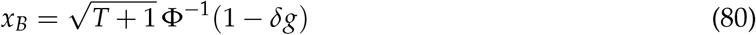

Since it is more convenient to have a constant bound, we choose *t* = *T* (Eq. 62) which provides us with the most conservative estimate.

##### 3.2.2 Time scale of the FPE

We are already measuring time in units of *t*_*g*_, such that it is adimensional. However, *T* still presents a large range of variation, so when solving the FPE it is convenient to measure time in units of *T*, such that it always goes from 0 to 1. In addition, if we define the *x* scale as 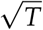, the FPE becomes^123^

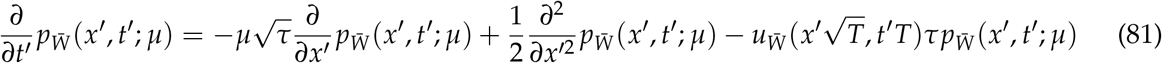

where 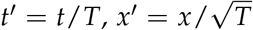. Therefore, to convert to real time, we multiply *t*^′^ by *t*_*g*_*T* (to undo both this latter change of variables and the initial one).

#### 3.3 Evaluation of log[*h*(*g*)]

The Bellman Eq. or the equations for the optimal policy contain terms of the form *h*(*g*) exp[−*βv*(*g, t*)], which we evaluate as

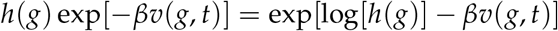

The quantity log[*h*(*g*)] can be numerically challenging. To avoid numerical instabilities, we write

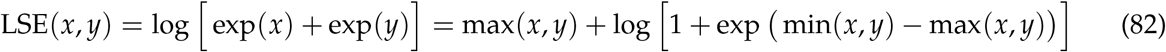

because the function log1pexp ≡ log[1 + exp(*x*)] can be implemented achieving high numerical precision for *x* ≤ 0. Then we can use the identities

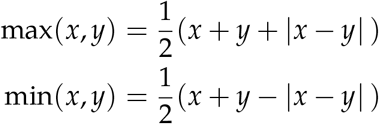

to write

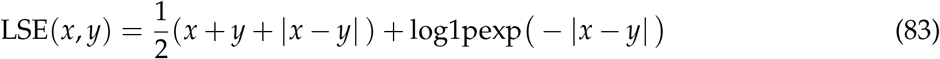

### 4 Sequential dependencies across trials

Although history effects generate long-timescale dependencies, where parameters in a particular trial might depend on events many trials in the past, their effect on the value function and optimal policy is tractable. This is, first, because in the average adjusted reward framework different trials only interact through the value of the reward rate, and second, because the effect of cross-trial dynamics on parameters only affects the reward rate through the steady state distribution of parameter values, as long as the dynamics of the parameters can be approximated as an ergodic Markov chain.

As described in the text, we consider sequential dependencies induced by across-trial dynamics in three parameters: a bias *ϕ* in the default policy (Eq. 26), the reward associated to correct responses to one of the options (which we will refer to as *r*), and the prior belief of the agent *g*_0_ at the beginning of the trial. Globally, we refer to these are bias-inducing-parameters (BIPs). We define the vector **X** as containing all sets of BIPs that might be encountered in a session

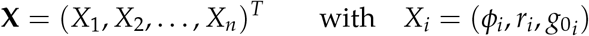

Since the Bellman Equation for the whole session in the average adjusted reward framework is identical to the one for a single trial except for the specification and interpretation of the reward rate *ρ* (section 1.4), for any *X*_*i*_ we can calculate the associated value function *V*_*i*_(*t, g*; *X*_*i*_), for a given *ρ*. One may interpret the *X*_*i*_ as extending the state space, which now includes the BIPs. The stimulus onset (i.e, *t* = 0) still corresponds to a proper reference state (section 1.4). Thus, to determine the value of the reward rate *ρ*, we define a mixture of the *V*_*i*_(*t, g*; *X*_*i*_) weighted by the stationary probability *π*_*S*_(*X*_*i*_), and use the condition that the value *V*_0_ of the mixture at *t* = 0 should be equal to zero (section 1.4), i.e.

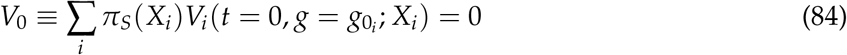

We find *ρ* using a similar iterative procedure as for the case without sequential dependencies. Since there is a single, constant *ρ* regardless of how many states *n* are part of **X**, at each step all *V*_*i*_(*X*_*i*_) are computed using the same tentative value of *ρ* to then re-evaluate Eq. (84) until convergence. Once the correct value of *ρ* has been found, then each *V*_*i*_ is re-evaluated, and we can directly compute the policies for each *X*_*i*_, which then allow deriving behavioral predictions (such as psychometric functions) for each BIP set.

In practice, to obtain the results in Figure 8, for each type of bias we consider BIP vectors of three states where only the correspondent parameter was changed in opposite directions, while the others were kept at their baselines:

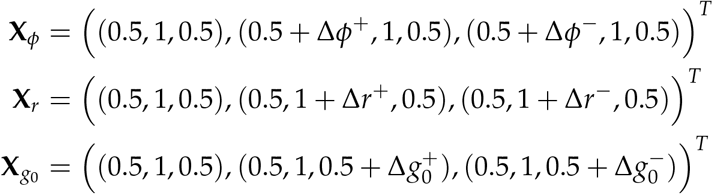

The stationary probability distributions were in all cases *π*_*S*_ = (1/3, 1/3, 1/3)^*T*^. The results do not change qualitatively for any reasonable choice of these probabilities. The procedure we just described is similar in spirit to a recently published method to categorize discrete behavioral states^24^. In fact, one can also include the parameters *β* and *λ* inside **X** and describe changes in control capability and responsiveness across trials. This should be addressed in future work.

#### 4.1 Biased passive policy

The passive policy is now

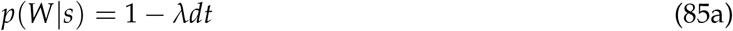

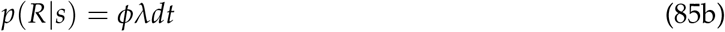

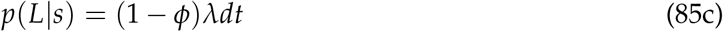

such that *ϕ* represents the bias towards responding *R*. The magnitude of *ϕ* enters the value function through a modification of *h*(*g*) in Eq. 57, which is now given by

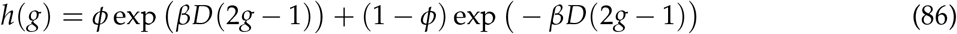

Given the new form for *h*(*g*) value function, policies and behavioral predictions proceed as before.

#### 4.2 Biased passive policy with biased rewards

We consider now a payoff matrix of the type

Payoff matrix(ℛ(*s, a*)):

|  | R | L | W |
| --- | --- | --- | --- |
| RS | $r$ | 0 | 0 |
| LS | 0 | 1 | 0 |

This causes the action value 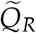 to be rewritten as

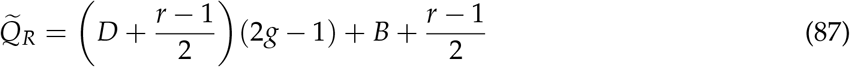

Therefore the function *h*(*g*) now is

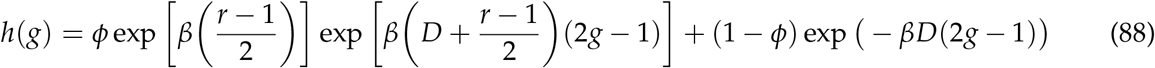

from which again value function, policies and behavioral predictions proceed as before.

#### 4.3 Biased prior beliefs

We implement the bias in the agent’s prior belief about the stimulus as a shift in the distribution *p*_0_(*μ*), which is effectively equivalent to a change in *g*_0_(*t* = 0), following Equation 29. Due to the one-to-one mapping between *g* and *x*, a positive (negative) value of *g*_0_ corresponds to a positive (negative) offset in the starting point of the evidence accumulation. Since the evolution of belief is unaffected by this change, none of the relevant equations need to be modified. One only needs to make sure that the value of the reference state used to calculate *ρ* is *V*(*t* = 0, *g*_0_), and that the initial condition of the FPE (Eq. 73) is shifted accordingly, producing the observed bias in the psychometric functions.

